# Granger-causal inference of the lamellipodial actin regulator hierarchy by live cell imaging without perturbation

**DOI:** 10.1101/2021.05.21.445144

**Authors:** Jungsik Noh, Tadamoto Isogai, Joseph Chi, Kushal Bhatt, Gaudenz Danuser

## Abstract

Many cell regulatory systems implicate nonlinearity and redundancy among components. The regulatory network governing lamellipodial and lamellar actin structures is prototypical of such a system, containing tens of actin-nucleating and -modulating molecules with functional overlap and feedback loops. Due to instantaneous and long-term compensation, phenotyping the system response to perturbation provides limited information on the roles the targeted component plays in the unperturbed system. Accordingly, how individual actin regulators contribute to lamellipodial dynamics remains ambiguous. Here, we present a perturbation-free reconstruction of cause-effect relations among actin regulators by applying Granger-causal inference to constitutive image fluctuations that indicate regulator recruitment as a proxy for activity. Our analysis identifies distinct zones of actin regulator activation and of causal effects on filament assembly and delineates actin-dependent and actin-independent regulator roles in controlling edge motion. We propose that edge motion is driven by assembly of two independently operating actin filament systems. A record of this paper’s Transparent Peer Review process is included in the Supplemental Information.

## Introduction

Many cell functions are regulated by biochemical and biophysical circuits with component redundancy, as well as feed-back and feed-forward interactions. A prototypical case of such a circuit is the machinery driving the formation of lamellipodia and lamella filamentous actin (F-actin) networks (Insall and Machesky, 2009, Isogai and Danuser, 2018). The dynamics of network assembly is controlled by dozens of actin-binding proteins with distinct structural and kinetic properties (Figure 1A) (Krause and Gautreau, 2014, Blanchoin et al., 2014, Insall and Machesky, 2009, Pollard, 2016, Pollard et al., 2000). The complexity in architectural dynamics is superimposed by complexity in biochemical signals, which orchestrate filament branching by the Arp2/3 complex, filament elongation by formins and Ena/VASP proteins, filament capping by Capping Protein (CP), and filament severing by ADF/cofilin proteins (not depicted) in response to cell-intrinsic and -extrinsic mechanical and chemical cues (Figure 1A). Although genetic and molecular perturbation has been instrumental in compiling an inventory of these system components and their basic contributions to lamellipodia formation, the functional hierarchy among the components has yet to be defined. In a system with so many functionally overlapping elements, perturbation of any one of them or any regulator upstream results almost instantaneously in a rewiring of the circuitry (Isogai and Danuser, 2018).

**Figure 1.**
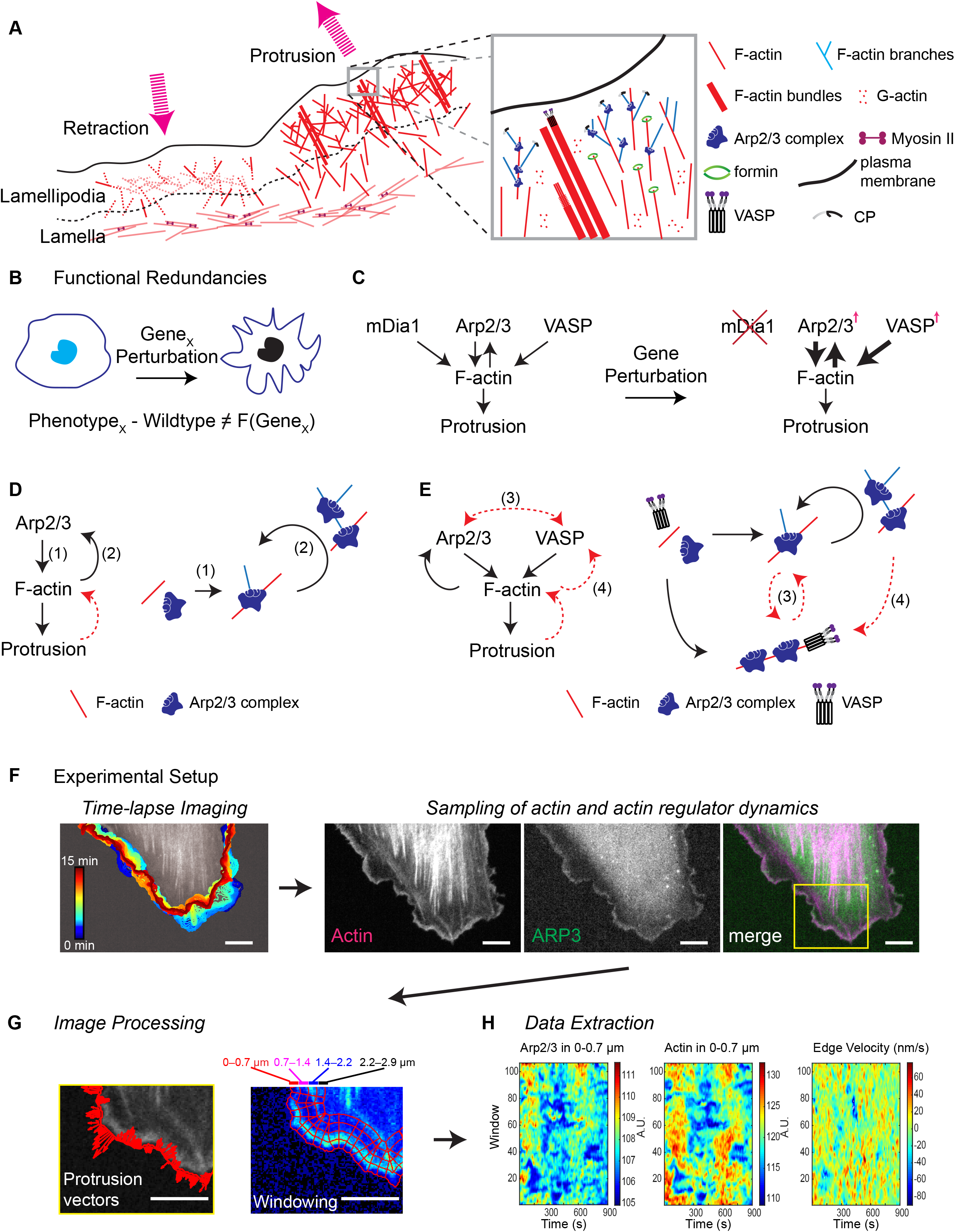
Actin network regulation in lamellipodia and lamella, a prototypical challenge in probing causal relations among molecular components. (A) Lamellipodia and lamella actin network formation at the edge of motile cells. Inset, select actin regulatory proteins that orchestrate the formation of branched, linear, and bundled actin structures. (B) In a perturbation experiment, the difference between phenotype and wildtype is non-equal to the function the targeted gene (Gene_X_) assumes in the unperturbed system. (C) In the system illustrated in (A) at least three proteins control the assembly of actin filaments (F-actin). Perturbation of any of these proteins will trigger compensation responses. (D, E) Examples of interactions between regulatory proteins and F-actin. Black arrows, well-established molecular interactions; these are used to validate the proposed causal inference framework. Red dotted arrows, hypothetical interactions with some rationale from in vitro biochemistry; these are tested using the proposed causal inference framework. (F) Imaging pipeline to acquire fluctuation time series of edge dynamics and molecular activities suitable for Granger-causal pathway inference. Proteins under scrutiny are fluorescently labeled and imaged over time. Scale bars, 5 µm. (G) Computer vision algorithms track cell boundaries and probing windows to extract cell edge protrusion/retraction velocities and spatiotemporal recruitments of molecules. Scale bars, 5 µm. (H) For each molecule, intensity fluctuation time series are extracted per window and visualized in a heatmap. See also Videos 1–3.

In the face of redundancy and nonlinearity, phenotypes induced by experimental perturbation are in a strict sense uninterpretable with respect to the role the targeted component plays in the unperturbed system (Vilela and Danuser, 2011, Welf and Danuser, 2014). Phenotypes show how the system adapts to the intervention, which is unequal to the function the targeted component assumes in the intact system (Figure 1B). For example, the linear F-actin nucleator formin mDia1, the branched actin nucleator Arp2/3 and the filament elongator VASP collectively contribute to the formation of actin networks, which in turn drives membrane protrusion (Insall and Machesky, 2009, Krause and Gautreau, 2014, Blanchoin et al., 2014, Pollard et al., 2000, Goode and Eck, 2007, Krause et al., 2003, Courtemanche, 2018). Perturbation of mDia1 will affect the actin network and subsequent membrane protrusion, but the phenotype, if observable, is confounded by altered activation of Arp2/3 and VASP (Figure 1C). Of note, the disconnect between phenotype and component function is intrinsic to the circuit wiring and does not relate to the widely-discussed additional complication of cellular adaptation to long-term perturbation (Hoeller et al., 2014). Moreover, feedback relations often escape conventional perturbation approaches. For example, Arp2/3 and F-actin are in a forward relation Arp2/3 → F-actin (Figure 1D(1)), but based on extensive biochemical experimentation there may be also a positive feedback F-actin → Arp2/3 (Figure 1D(2)) (Mueller et al., 2017, Bieling et al., 2016). How important is this feedback eventually for the growth of the branched lamellipodial actin network? Answering this question is non-trivial and gets more complicated when considering the concurrent contributions of additional factors such as VASP. VASP accelerates elongation of pre-formed filaments. Will the elongated filaments recruit more Arp2/3 and in turn more F-actin nucleation (Figure 1E(3))? And will the newly nucleated F-actin serve as an additional substrate for VASP (Figure 1E(4))? To dissect circuitry of such complexity the field needs novel approaches that overcome the limitations of probing by perturbation.

**Figure 2.**
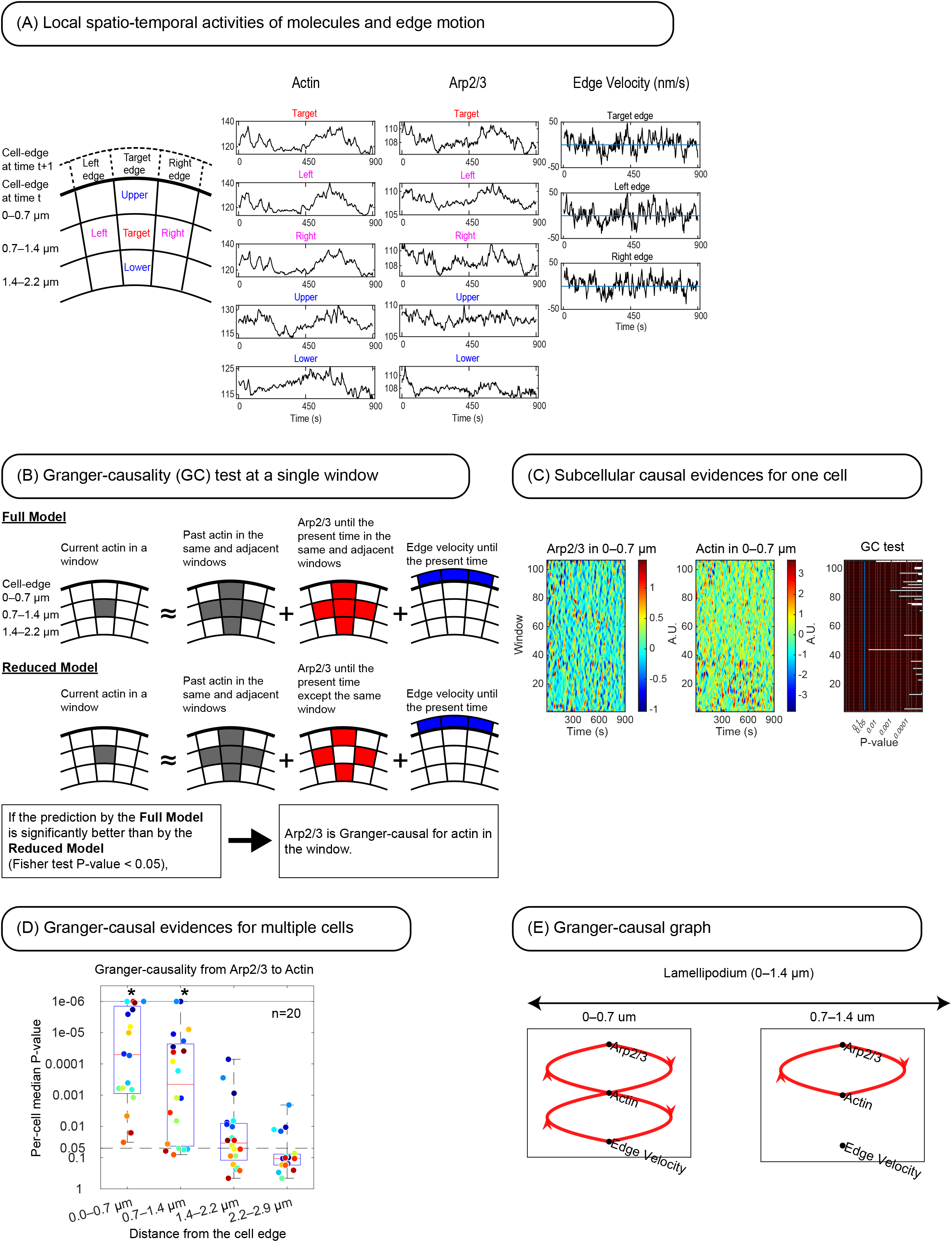
Workflow of Granger-causal pathway inference in the study of lamellipodial actin dynamics. (A) Example of actin and Arp2/3 recruitment time series sampled in the target window and its immediate neighbors. For windows at the cell edge time series of edge motion are sampled. (B) A full regression model specifies the relation between actin recruitment to the target window at time *t* and the time series of all variables shown in (A) up to the time point t. A reduced regression model specifies the same relations except the Arp2/3 recruitment to the target window. The Granger-causality (GC) test determines whether the full model predicts the actin activities significantly better than the reduced model due to essentiality of the omitted information of Arp2/3 recruitment. (C) P-values of the GC tests in individual windows provide subcellular evidence of the causal link from Arp2/3 recruitment to F-actin assembly. Shown are windows of layer 1 (0 – 0.7 μm from the cell edge). (D) The subcellular GC P-values for a particular window layer are integrated into a per-cell median P-value for that layer. Per-cell median values are aggregated in a distribution and tested for the median of the distribution < 0.05 based on the Wilcoxon signed rank test. (E) Significant Granger-causal relations are represented as graphs, drawn separately for probing window layers at increasing distances from the edge.

**Figure 3.**
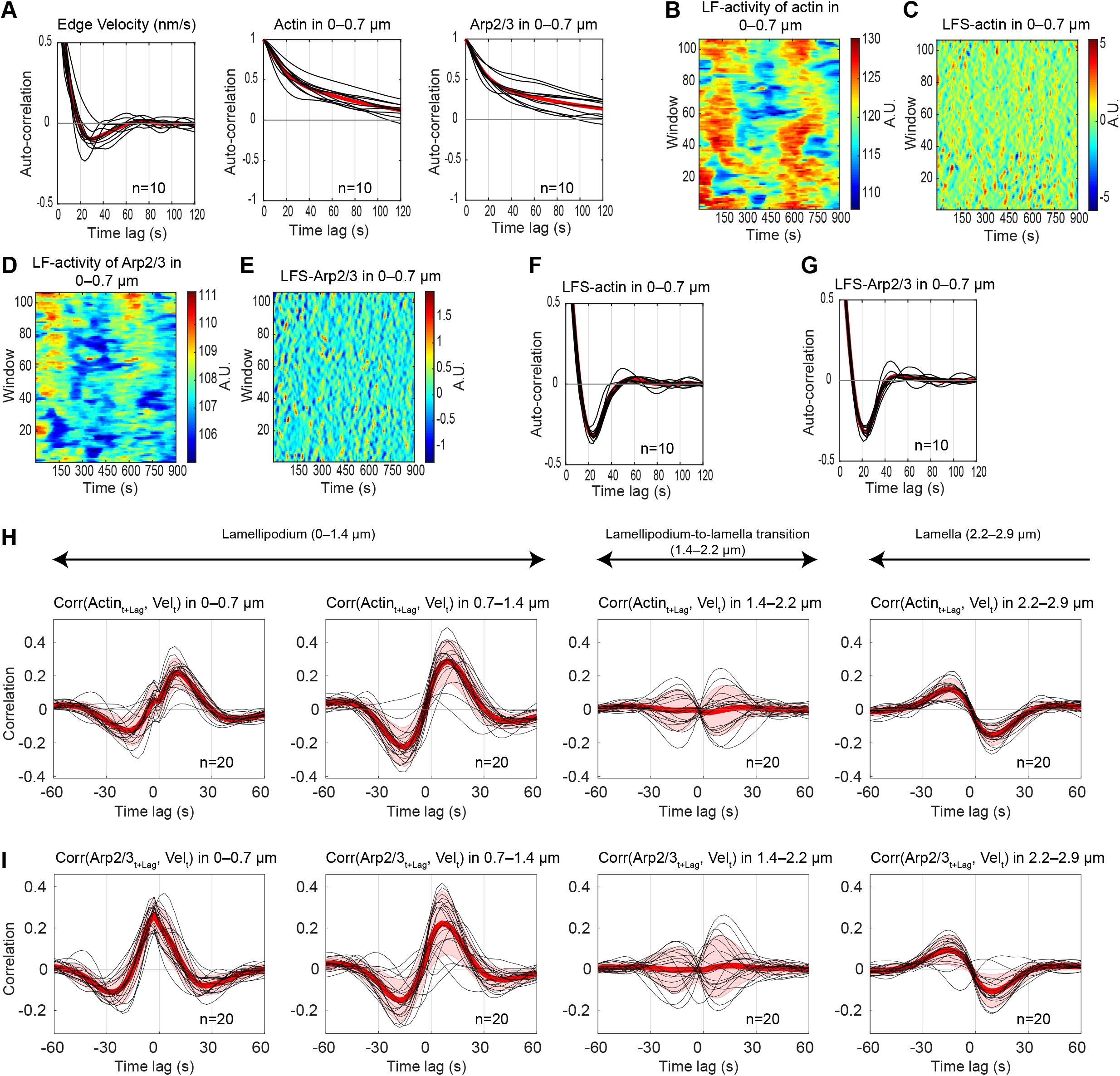
Actin and Arp2/3 intensity fluctuations correlate with edge motion. (A) Auto-correlation functions (ACFs) of edge velocity and the recruitment of actin and Arp2/3 in layer 1 (0–0.7 µm from the cell edge). Black curves, per-cell averaged ACFs; red curves, population average (n = 10 cells). (B-C) Spatiotemporal maps of low-frequency (LF) fluctuations (B) and low-frequency subtracted (LFS) fluctuations (C) of actin assembly in layer 1. (D-E) LF fluctuations (D) and LFS fluctuations (E) of Arp2/3 recruitment in layer 1. (F-G) ACFs of LFS-recruitment of actin (F) and Arp2/3 (G) in in layer 1. Black curves, per-cell averaged ACFs; red curves, population average (n = 10 cells). (H-I) Per-cell averaged cross-correlation curves (n = 20, black) of LFS-actin (J) and LFS-Arp2/3 (K) with the edge velocity. Red curves, population averages; shaded bands, ±2 × SEM. See also Video 4.

**Figure 4.**
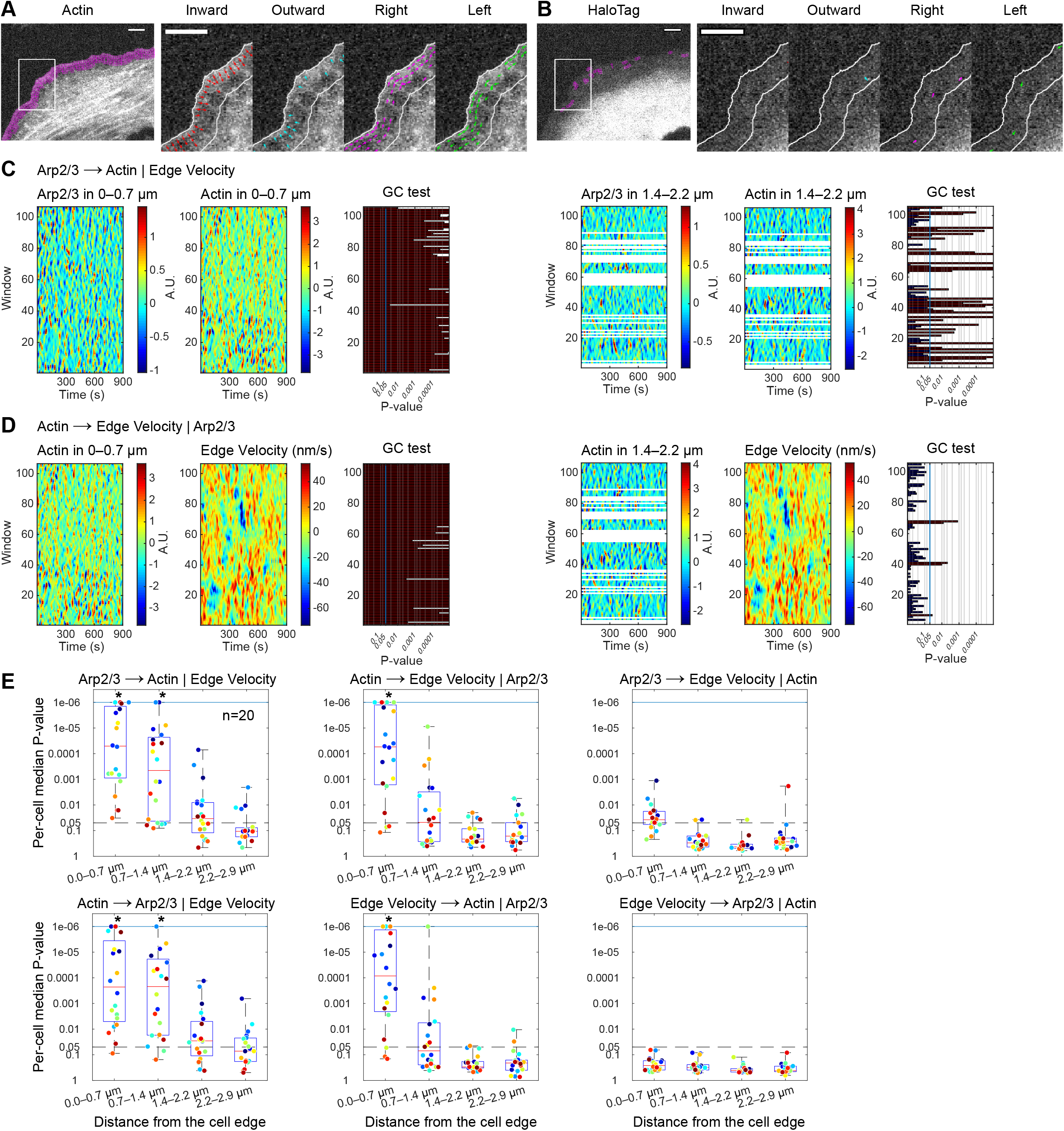
Granger-causality analysis establishes a causal chain from Arp2/3 to F-actin to edge motion. (A-B) Spatial propagation patterns of molecular activities annotated on a single time point image of a U2OS cell co-expressing mNeonGreen-tagged actin (A) and cytoplasmic HaloTag (B). Magenta windows (left panel), subcellular regions where molecular fluctuations propagate to four adjacent windows. Arrows (right panel), spatial propagation in the four different directions. Scale bars, 5 µm. (C) Arp2/3 and actin activity maps and the associated P-values of GC tests from Arp2/3 to actin in individual windows for a representative cell. White regions in the map for layer 3 (1.4–2.2 µm from the cell edge), drop out of probing windows due to the convergent window grid topology away from the cell edge. Red, significant P-values. (D) GC tests from actin to the edge velocity, represented as in (C). (E) Distributions of per-cell median P-values for six directional Granger-causal relations between Arp2/3, actin and edge velocity at different distances from the cell edge (n = 20 cells). The symbol (*) indicates per-cell median P-values significantly smaller than the nominal level 0.05 (Wilcoxon signed rank test). See also Videos 5–12.

Several studies of actin regulation during cell protrusion and retraction have exploited live cell fluorescence time-lapse imaging and computer vision as an observational approach to infer the underlying molecular pathway structure without experimental intervention (Mueller et al., 2017, Bisaria et al., 2020, Barnhart et al., 2017, Lee et al., 2015, Welf and Danuser, 2014, Isogai and Danuser, 2018, Ji et al., 2008). Fluorescence intensity fluctuations reporting the recruitment of actin modulators to the F-actin network as a surrogate of their activity were extracted from live cell movies. Fluctuation time series were then exploited to establish the correlation between molecular and morphological activities, assuming that the numerical coupling of the two is an indicator of local functional relations (Ponti et al., 2004, Lee et al., 2015, Ji et al., 2008, Mendoza et al., 2015, Mueller et al., 2017, Wang et al., 2018, Bisaria et al., 2020). This paradigm has also been applied to the analysis of regulatory signals upstream of actin dynamics (Machacek et al., 2009, Azoitei et al., 2019, Yang et al., 2016, Marston et al., 2020). However, these analyses do not inform on the causal relation between activities, especially not in the face of feedback relations.

Given a set of temporally resolved variables, the hierarchy of cause-and-effect relations can be inferred by statistical assessment of the power of the signal of a putative cause for the prediction of the signal of a putative effector. This notion of inferring causality has long been employed in econometrics and neurophysiology (Granger, 1969, Bressler and Seth, 2011). The most popular of these frameworks is the Granger-causality (GC) analysis, which defines a statistical test of the hypothesis that past observations of one variable possess indispensable information for explaining the current and future observations of a second variable (Eichler, 2013). Because of the explicit temporal direction in the relationships, the GC framework also permits analysis of nonlinear regulatory motifs such as feedback, redundant pathways, and even nested feedbacks.

Granger-causal (G-causal) relations must be interpreted only within the system of co-observed variables. For example, if the observed signal of a variable X is causative for an unobserved latent factor that is causative for the observed signal of a variable Y, then X will be determined as G-causal for Y. While in many biological studies, knowledge of such indirect relations can yield great insight for practical purposes, the prediction of a G-causal relation is not to be mistaken for a causality that pinpoints direct molecular interactions. Accordingly, we refer to G-causal relations as indicators of functional causality. In contrast, the prediction of Granger-noncausality has the strong implication that the two considered variables are independent, regardless of any latent factor. This property permits the exclusion of functional relations at the level of the whole system based on a partially observed system (Eichler, 2007).

In this work, we illustrate the potential of GC analysis in overcoming the challenges of deciphering the molecularly entangled regulation of lamellipodial F-actin assembly. Using multivariate time-series representing the spatiotemporal molecular and cell morphological dynamics of the system, we build generative stochastic models that capture the causal structure of the system. Our GC analytical pipeline, for example, shows that actin dynamics at the most proximal zone, i.e. ∼0–0.7 µm from the cell edge, Granger-causes (G-causes) edge motion, while actin more distal from the cell edge (zones ∼0.7–1.4 µm) is predicted as Granger-noncausal (G-noncausal) for edge motion. Notably, this latter zone showed a strong positive correlation between F-actin dynamics and edge motion, demonstrating that GC analysis is distinct from correlation analysis: Correlation is not causation. We further applied the GC pipeline to identify the causal relations between actin modulators and F-actin dynamics. For Arp2/3 and VASP, for example, our analysis determines that both modulators G-cause edge motion yet operate independently from each other – a conclusion that has been inaccessible by conventional perturbation and correlation-dependent approaches.

## Results

### Extraction of image fluctuation time series of actin regulator recruitment suitable for Granger-causality analysis

To study G-causal relations among actin regulators, F-actin assembly and lamellipodia dynamics, we acquired fluorescence time-lapse images of cells expressing either endogenously tagged or low concentration of tagged ectopic regulator protein, so as to not perturb the endogenous stoichiometry among regulators. By ensuring low expression of the labelled regulator (see Methods), local changes in fluorescence intensity largely reflect the association of the regulator with F-actin (Danuser and Waterman-Storer, 2006). We illustrate the time series extraction based on the example of U2OS cells co-expressing Arp3 labelled with CRISPR/Cas9-edited C-terminal HaloTag (Halo) and mNeonGreen-tagged actin under a truncated CMV promoter. Cycles of edge protrusion and retraction events were sampled every 3 seconds for 15 minutes (Figure 1F). Visual inspection of these cycles in conjunction with F-actin dynamics and Arp3 recruitment revealed the well-established characteristics of a lamellipodium with actin retrograde treadmilling at the cell front (∼1.5 µm from the edge) and a lamella with slower and spatially less coherent actin dynamics beneath (>1.5 µm from the edge) (Ponti et al., 2004) (Video S1).

To record fluctuation time series of edge motion and cytoskeletal dynamics, we partitioned the protruding and retracting front of the cell into submicron-scale probing windows (Figure 1G), and tracked their positions over time so that they maintained a constant relation with an edge sector of the same submicron scale, as described previously (Ma et al., 2018, Machacek et al., 2009). This generates a coordinate system that allows simultaneous registration of spatiotemporal fluorescent intensity fluctuations and cell edge protrusion/retraction dynamics (Video S2-S3). The size of the probing windows was fine-tuned to be several-fold smaller than the average length scale of protrusion and retraction events, and small enough to capture the distinct cytoskeleton dynamics in lamellipodium and lamella. Specifically, the lamellipodia depth in U2OS cells measured 1.4 µm, on average. Thus, for our spatial analyses, we defined the lamellipodia region as the band ∼0–1.4 µm from the cell edge and the lamella region as the band ∼1.4–2.9 µm from the cell edge. We further divided these regions into half to examine potential spatial gradients in Arp2/3 and actin activities within each region. For each probing window we then read out time series of locally averaged Arp2/3 and actin intensities and mapped them into space-time matrices (Figure 1H; see Methods). For the window row at the cell front, we also read out the average velocity, with positive and negative values indicating protrusion and retraction, respectively. Of note, any probing window in layer 2 and higher is unambiguously associated with one probing window in layer 1. This permits the analysis of causal relations between different types of events, for example, Arp2/3 and velocity, or Arp2/3 and another actin regulator, in the same or different window layers.

### Workflow of Granger-causal pathway inference between regulators of lamellipodial actin dynamics

To illustrate the inference of G-causal relations we focus on the questions ‘how causal’ is Arp2/3 recruitment to a particular target window for the assembly of actin in the same window, and ‘how causal’ is the assembly for cell edge motion at that location. Common expectation in the field would suggest the existence of a causal chain Arp2/3 → F-actin → motion. Thus, we can use the data in Figure 1H to validate the basic application of GC-analysis for this system. However, even with these extremely well-studied relations, it is unknown at what distance from the cell edge they may break and whether there is feedback from F-actin to Arp2/3 recruitment (Figure 1D). Both questions can be uniquely addressed by GC-analysis.

Per Granger’s definition (Granger, 1969), Arp2/3 recruitment is G-causal for F-actin, if the best prediction of the F-actin assembly time series using ‘all available information’, including the Arp2/3 recruitment, is significantly better than the best prediction of F-actin assembly using all information except the Arp2/3 recruitment. Hence, in a particular target window we need to assess whether knowledge of the Arp2/3 recruitment supports the prediction of F-actin assembly in the same window. The F-actin assembly time series in the target window may further be influenced by the Arp2/3 and F-actin recruitment in the neighboring probing windows. Indeed, the high level of similarity between fluctuation time series in windows surrounding the target window suggest a strong biochemical coupling of the actin regulatory process in space (Figure 2A). To determine the G-causal relation between Arp2/3 and F-actin in the target window, contributions from time series in the surrounding windows must be considered as confounding factors. The model also accounts for the protrusion/retraction (P/R) velocities as potential confounders, i.e. the analysis considers whether Arp2/3 and F-actin co-fluctuate simply because they jointly follow the edge movements. Taken together, the indispensability of Arp2/3 recruitment for the prediction of F-actin assembly at a given location is tested by comparing two models (Figure 2B, see Methods): (1) the full model incorporating fluctuation time series of Arp2/3, F-actin and associated edge motion in the target window and the four neighboring windows (Methods Eq. 2), and (2) the reduced model equivalent to the full model minus the Arp2/3 fluctuation time series in the target window. If Arp2/3 is an indispensable variable, the full model will lead to a significantly better prediction performance than the reduced model. Because of the additional degrees of freedom, the full model will always perform better than the reduced model, as assessed by the variance of the difference between predicted and measured F-actin fluctuations. The key question is whether the additional degrees of freedom *significantly* improve the prediction. This is answered by application of a Fisher test under the null-hypothesis that the full and reduced models are equally strong predictors (Methods Eq. 3). In the context of Figure 2, low P-values, therefore, indicate a causal link from Arp2/3 recruitment to F-actin assembly.

G-causal links describe direct interactions between the observed variables. For example, in a pathway A → B → C A is referred to as directly causal for B, and indirectly causal for C. Hence, the G-causal link from Arp2/3 recruitment to F-actin assembly cannot be mediated by edge motion. If the pathway were to consist of a chain Arp2/3 → edge motion → actin, then edge motion would have accounted for F-actin assembly in the full and reduced models, rendering Arp2/3 dispensable for the prediction.

To draw firm and reproducible conclusions on the causality between variables, we integrated the subcellular P-values of the GC tests over multiple, independently imaged cells. From a statistical perspective, the *sampling unit*, i.e. the physical entity that is repeatedly and independently measured in our imaging experiments, is one cell. Because a statistical conclusion is a statement on the representative property of a population of sampling units, hypotheses on causal relations need to be tested based on per-cell observations. To accomplish this, we computed the median of P-values of GC evidences over the probing windows of a cell as the per-cell observation of causality. A per-cell median P-value of less than 0.05 indicates that the majority of probing windows in the cell show G-causal interactions between the investigated variables (Figure 2C). Subsequently, we apply the Wilcoxon signed rank test to determine whether the per-cell median P-values of *n* cells are less than 0.05, in which case the two investigated variables are considered as causally connected (Figure 2D).

After completing the pairwise testing of G-causal relationships between all variables in the observed system, causal relations are integrated and represented as a graph (Sugiyama et al., 1981) (Figure 2E). To account for spatial variation in causal interactions, we compute graphs separately for lamellipodia and lamella (in our U2OS cell model: 0–0.7 µm and 0.7–1.4 µm vs 1.4–2.2 µm and 2.2–2.9 µm, respectively). For the example of Arp2/3, F-actin, and edge velocity, the graphs indicate a causal interaction between Arp2/3 and actin, as expected (Pollard et al., 2000, Isogai et al., 2015, Goley and Welch, 2006, Blanchoin et al., 2000a), as well as a feedback from F-actin to Arp2/3, demonstrated for the first time in living cells. This feedback may be explained by the intrinsic recruitment dynamics of the dendritic polymer network, in which the nucleation of branches yields additional filaments that in turn can be branched again. Our analysis further shows that actin is in a causal feedback relation with edge velocity, however only for the probing windows in layer 0–0.7 µm. The forward link from actin to edge motion describes the well-established conversion of actin filament growth into mechanical push of the cell edge through mechanisms such as the Brownian Ratchet (Mogilner and Oster, 1996). The feedback from edge motion to F-actin assembly may be governed by several mechanisms including mechanical and chemical force-feedback from the increasing membrane tension or membrane deformation (Mueller et al., 2017, Higashida et al., 2013, Batchelder et al., 2011, Bieling et al., 2016, Lou et al., 2019). Importantly, the graphs indicate that there is no direct causal interaction between Arp2/3 and velocity. This means that any modulation in Arp2/3 recruitment translates into modulation of edge velocity only via F-actin dynamics. The absence of a causal link from velocity to Arp2/3 implies that the predicted feedback from edge motion to actin in layer 0–0.7 µm is independent of Arp2/3, i.e. signals must exist that translate the morphological dynamics or variation in membrane tension into actin nucleators other than Arp2/3. Hence, this data refutes the proposal that the bending of actin filaments under plasma membrane load contributes significantly to Arp2/3 recruitment (Risca et al., 2012). These first results demonstrate how Granger-causality analyses functionally relate molecular processes in a hierarchical and nonlinear order.

### Actin and Arp2/3 fluctuations cross-correlate with edge velocity, but so do fluctuations of a cytosolic tag

We also present the cross-correlations (CCs) between Arp2/3, F-actin, and edge velocity. For both CC- and GC-analyses it is essential that the time scales of variable fluctuations match. A straightforward approach to determine the time scale of stochastic time series relies on the auto-correlation function (ACF), which for protrusion and retraction projects average cycles of ∼60 s but cycles of > 4 min for F-actin and Arp2/3 recruitment (Figure 3A).

Since Arp2/3 and F-actin dynamics are known to be related to protrusion and retraction dynamics (Pollard et al., 2000, Krause and Gautreau, 2014), we hypothesized that the overall F-actin and Arp2/3 fluctuation signals contained shorter time scale fluctuations that would align with that of edge motion cycles whereas the dominant cycles at longer time scales are unrelated to cell morphodynamics. To test this, we decomposed the raw time series of F-actin and Arp2/3 recruitment into low-frequency (LF) oscillations and low-frequency subtracted (LFS) oscillations (Figures 3B-3E, see Methods). Indeed, the ACFs of the LFS signals closely matched the ACF of edge velocities (Figures 3F and 3G, Video S4). Hereafter, we refer to LFS-molecular activity without specifying the LFS for brevity.

We computed the cross-correlation (CC) between LFS-actin or -Arp2/3 and edge velocity in each window as previously described (Machacek et al., 2009). Within the lamellipodia (layers 1 and 2; ∼0–1.4 µm), the fluctuations of F-actin assembly best correlated with the edge velocity with a time delay of ∼9 s (Figure 3H), which is in line with previous reports (Ji et al., 2008, Lee et al., 2015). In contrast, Arp2/3 fluctuations preceded edge velocity fluctuations by ∼3 s in layer 1, whereas in layer 2, Arp2/3 followed the edge velocity by ∼6 s (Figure 3I).

Our correlation and GC-analyses operate under the assumption that intensity fluctuations in F-actin and Arp2/3 faithfully report the addition or removal of actin subunits and Arp2/3 complexes to or from the lamellipodial network. To test the validity of this assumption, we performed correlation analyses with images of diffuse HaloTag alone. The CC curves between cytoplasmic HaloTag intensities and edge velocity consistently showed positive correlation values at negative time lags and negative correlations at positive time-lags (Figure S1A). To explain the mechanism underlying the positive and negative lobes we turned to kinetic maps, which indicate the average accumulation of a fluorescent signal relative to a morphodynamic event such as protrusion or retraction onset (Azoitei et al., 2019, Lee et al., 2015) (Figure S1B). The typical fluorescence intensities of the cytoplasmic HaloTag reached its highest values after the fastest retraction, which explains the negative correlations between the HaloTag and edge velocity at positive time-lags. This peak of the HaloTag mean intensity was still observable before the fastest protrusion, which was captured by the positive correlations at negative time-lags. These fluctuation patterns of cytoplasmic HaloTag depicted systematic volume changes near the edge during P/R cycles, where the volume was maximized right before the protrusion onset and minimized at or right before the retraction onset as the cell edge maximally stretched out.

The CC pattern of a diffuse, cytoplasmic HaloTag with edge velocity resembled the CC patterns of both actin and Arp2/3 with edge velocity in the most distal region we analyzed (layer 4; 2.2–2.9 µm) (Figures 3H and 3I). This suggests that in the lamella, the LFS F-actin and Arp2/3 signals reflect a fraction of fluorescent probes that are diffusing in contrast to the relatively static cytoskeleton structures such as cortical actin fibers, whose fluorescence fluctuations are captured by the LF baseline recruitment signal. The CC patterns of F-actin and Arp2/3 recruitments with edge velocity in the transitional region 1.4–2.2 µm (layer 3) exhibited a mixture of the lamellipodial and cytoplasmic CC curves. This shows that the CC curves of F-actin and Arp2/3 differentiate two distinct cytoskeleton behaviors in lamellipodium and lamella, and that the lamellipodium-to-lamella transition in U2OS cells locates in the 1.4–2.2 µm region.

### Beyond correlation – a generative stochastic model to capture spatial information flows

Since cytoplasmic signals still correlated with edge velocity, correlations alone are insufficient to identify causal relations among the molecules involved in lamellipodial dynamics. To accomplish causal inference, we had to formulate a generative, stochastic model that specifies the dependencies of the edge velocities on molecular activities in subcellular space and time. Towards such a model, we first assembled a spatially propagating autoregressive (SPAR) model that prescribes the propagation of molecular activities in a target window to four adjacent windows (Figure S1C, see Methods). Analysis of actin dynamics by the SPAR model detected the propagation of F-actin signal fluctuations in retrograde direction in regions with prominent actin retrograde flow, as expected (Figure 4A, Video S5). It also predicted that actin fluctuations propagate laterally throughout the entire lamellipodia. In contrast, per the SPAR model fluctuations in the cytoplasmic HaloTag signal did not spatially propagate in most lamellipodia/lamella regions (Figure 4B, Video S6). This illustrates the capacity of SPAR analysis to differentiate between spatiotemporally propagated and non-propagated molecular activities.

### GC analysis identifies a causal chain from Arp2/3 to F-actin to edge velocity at the lamellipodia front

We extended the SPAR model to capture the relations between two spatiotemporally coupled molecular activities and edge velocity (Figure 2B). Applied to the analysis of lamellipodia dynamics, we first tested the causal effect of Arp2/3 recruitment on F-actin assembly at each subcellular probing window in the cell shown in Figure 1F. Most windows in the lamellipodia region (layers 1 and 2; ∼0–1.4 µm) displayed significant GC P-values, whereas the Arp2/3 did not G-cause actin fluctuation in the majority of windows outside the lamellipodia (layers 3; ∼1.4–2.2 µm) (Figures 4C and S2A). As one would expect, F-actin fluctuations at the lamellipodia front (layer 1) had G-causal effects on edge motion, with almost all probing windows showing P-values < 0.0001 (Figure 4D). The G-causal effect sharply dropped to a median P-value of 0.032 over all windows at the lamellipodia base (layer 2, Figure S2B). Beyond the lamellipodium, F-actin fluctuations stayed G-noncausal for edge velocity in most windows (median P-values 0.419 and 0.571 over all windows in layers 3 and 4, respectively; Figure 4D and not shown). For the remaining combinations of pairwise relations between F-actin, Arp2/3 and edge velocity, we found distinct subcellular patterns of causality reflecting the fine-grained spatial regulation of the lamellipodial and lamellar actin machinery (Video S7-S12). Together, these analyses demonstrate the high spatial resolution of the SPAR models in delineating spatial zones of distinct causal hierarchy among subcellular events.

We then examined whether the patterns of G-causal relations identified in a single cell were reproducible in cells imaged over independent experimental sessions (Figure 4E). Applied to a population of n = 20 cells, these statistical analyses confirmed a causal relation from Arp2/3 to F-actin assembly in layers 1 and 2 (∼0–1.4 µm), as expected. Our data also identified that F-actin assembly G-causes Arp2/3 recruitment (rank test P-values < 0.001), confirming the proposal of an autocatalytic dendritic nucleation put forward in Figure 1D as one of the hypotheses difficult to be tested by intervention approaches (Pollard and Borisy, 2003, Blanchoin et al., 2000b, Carlsson, 2003, Isogai et al., 2015). Both forward and feedback causal relations between Arp2/3 and F-actin were insignificant in layers 3 and 4 (rank test P-values > 0.522, Figure 4E), demonstrating that Arp2/3-actin interactions as they relate to P/R cycles are confined to the lamellipodia area. Similarly, the analysis of median P-values in cell cohorts supported a strong G-causal relation from F-actin to edge motion in layer 1 (rank test P = 0.001, Figure 4E), but no significant relation deeper into the lamellipodia (rank test P = 0.935; Figure 4E).

A critical insight gained from a SPAR model that combines Arp2/3, F-actin, and edge motion is that Arp2/3 recruitment at the lamellipodia front is G-noncausal for edge velocity (rank test P = 0.135, Figure 4E), consistent with a causal cascade Arp2/3 → F-actin → edge velocity. Hence, any causal effect from Arp2/3 recruitment to edge motion is fully mediated by F-actin assembly. Indeed, when we excluded the F-actin data from the SPAR model, the GC pipeline predicted a strong G-causal relation from Arp2/3 to edge velocity in layer 1 (rank test P < 0.001), but not in layer 2 (rank test P = 0.977, Figures S2C-S2E). This shows that the mechanism of GC inference distinguishes direct from indirect causal effects among the observed components of a molecular system.

The change in GC topology between the two- and three-component systems (Figures S2F-S2H and 4E) also rules out the possibility that any other variable besides F-actin mediates the causal effect of Arp2/3 on edge velocity. If such a hidden mediator were to exist, then the GC analysis in the three-component model would predict an additional G-causal relation from Arp2/3 to edge velocity bypassing F-actin.

To further validate the specificity of GC inference, we applied the pipeline to an inert HaloTag cytoplasmic control data (n = 12 cells). As expected, the analysis determined that cytoplasmic HaloTag fluctuation is G-noncausal for F-actin assembly or edge velocity (Figure S3, rank test P-values > 0.633), although the fluctuations correlate with edge motion quite strongly (Figure S1A). This again underlines the ability of the presented pipeline to separate causal from correlative relations.

### mDia1 recruitment G-causes F-actin assembly at the lamellipodia base and in the lamella and plays an actin-independent role at the lamellipodia front

Using the GC pipeline we set out to determine where other actin nucleators and modulators affect actin assembly and cell edge protrusion/retraction. We started with the formin family member mDia1. Formins are known to bind to the barbed-ends and processively elongate linear actin filaments (Pollard, 2016, Courtemanche, 2018). We stably depleted endogenous mDia1 in U2OS cells using lentiviral short hairpin RNA and introduced exogenous SNAP-tagged mDia1 and mNeonGreen-tagged actin under a truncated CMV promoter to follow their dynamics. Live cell movies displayed dynamic recruitment of mDia1 near the cell edge along with the edge protrusion/retraction cycles (Figure S4A, Video S13). We found no G-causal interactions of mDia1 with either actin or edge motion (Figures S4B and S4C), which seemed to contradict previous reports in other cell types of mDia1 functioning as the initiator of actin assembly and protrusion (Isogai et al., 2015, Lee et al., 2015). We suspected that the level of overexpression of mDia1 (Figure 5A), albeit experimentally controlled as much as possible, resulted in a significant perturbation of the stoichiometry among the nucleators, masking mDia1’s proper function. We thus inserted by CRISPR/Cas9 a SNAP-tag before mDia1’s N-terminal sequence but found the resulting fluorescence movies to have too low signal-to-noise ratio (SNR) for a meaningful causality analysis (data not shown). To overcome the deficiency in SNR, we decided to endogenously tag mDia1 with a tandem mNeonGreen2-based split fluorescence protein (sFP) strategy (Kamiyama et al., 2016) (Figure 5B), which primarily suppresses background, yet also amplifies the signal.

**Figure 5.**
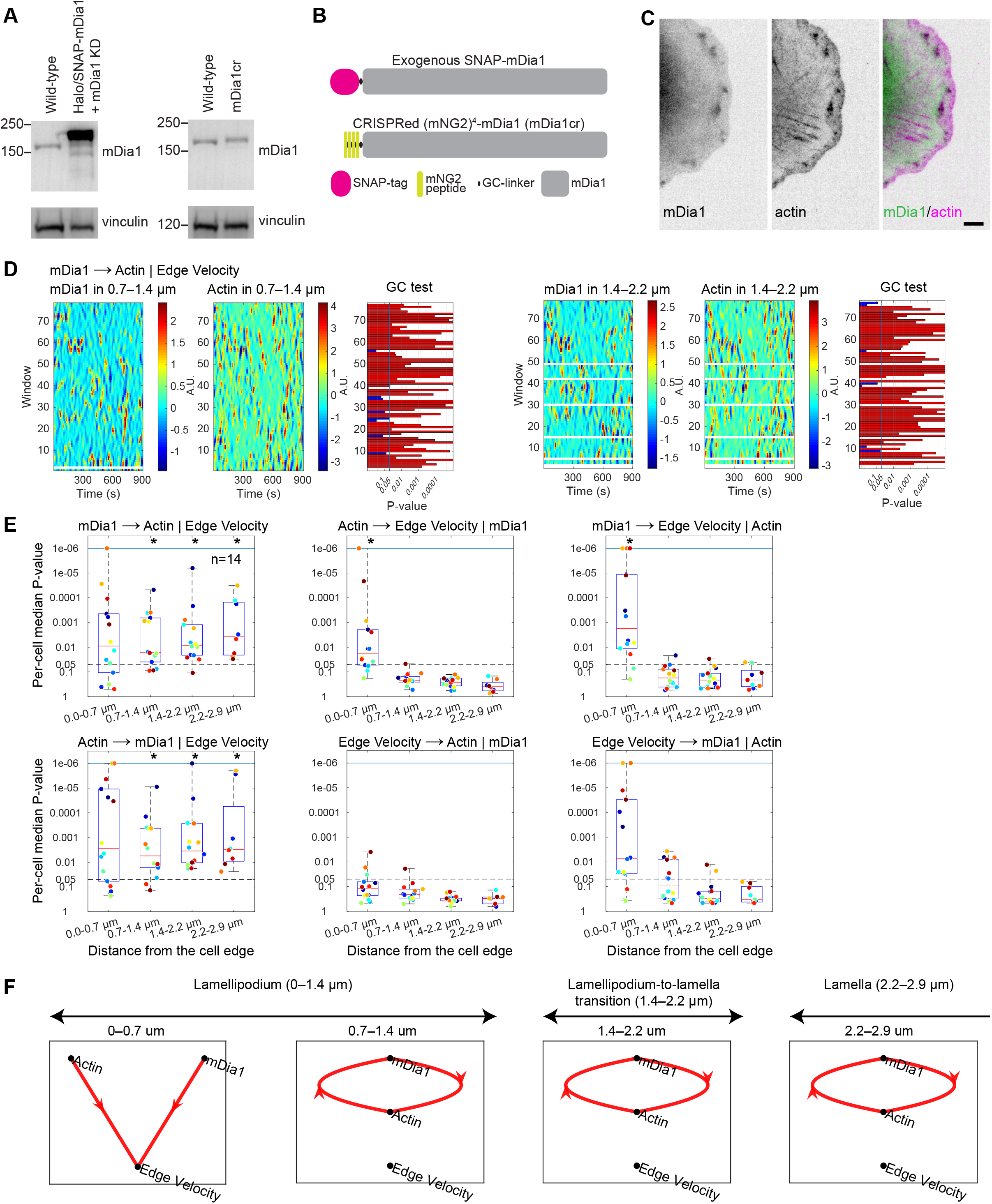
mDia1 fulfills spatially segregated, actin polymerization-dependent and polymerization-independent functions in the lamellipodia. (A) Expression of exogenous SNAP-tagged mDia1 (left panel) and endogenously labeled mNG_11_-mDia1 (right panel). (B) Comparison of molecular sizes of the tagged mDia1s drawn to scale. (C) Representative image of SNAP-actin and endogenously labeled mNeonGreen2-mDia1. Scale bars, 5 µm. (D) mDia1 and actin activity maps and associated P-values of GC tests from mDia1 to actin in the lamellipodia back (left panel) and in the lamellipodia-to-lamella transition area (right panel) for the cell in (C). Red, significant P-values. (E) Distributions of per-cell median P-values of Granger-causal relations between mDia1, actin and edge velocity (n = 14 cells). The symbol (*) indicates per-cell P-values significantly smaller than 0.05 (Wilcoxon signed rank test). (F) GC pathway diagrams between mDia1, actin and edge velocity in the lamellipodia and lamella regions. See also Video 14.

Live cell movies of endogenous mDia1 displayed multiple spots of dynamic recruitment a few microns back from the cell edge, which were unobservable with exogenous expression of tagged mDia1 (Figure 5C). The spots co-localized with spots enriched for actin and they appeared to be associated with strongly protruding cell edge segments (Video S14).

Using CC analysis, we found that, unlike F-actin and Arp2/3, mDia1 recruitment and edge velocity correlated homogenously across the lamellipodia and lamella. The CC curves over multiple cells (n = 14) consistently showed maximal negative values at positive lags of ∼3-9 s (Figure S4D). Kinetic maps indicated that this negative peak was associated with mDia1 recruitment after maximal retraction and before protrusion onset (Figure S4E), consistent with previous observations (Lee et al., 2015).

To determine how much mDia1 recruitment causes actin assembly and edge motion, we applied the GC pathway inference pipeline to mDia1/actin live cell movies. On a per-cell basis, most windows at the lamellipodia base and in the lamella (∼0.7–2.9 µm) showed significant G-causal effects of mDia1 on F-actin recruitment (Figures 5C and 5D). For the entire cell population (n = 14), the analysis determined causal influence of mDia1 on F-actin assembly at the lamellipodia base and lamella (∼0.7–2.9 µm, rank test P < 0.012, Figure 5E), but not at the lamellipodia front (0–0.7 µm, rank test P = 0.452). The observed G-causal feedback between actin on mDia1 (rank test P < 0.010, Figure 5E), may represent mDia1’s activity to recruit profilin-bound actin for F-actin nucleation and elongation.

Our analysis further identified a direct G-causal relation between mDia1 and edge velocity at the cell edge, where mDia1 is G-noncausal for F-actin assembly (0–0.7 µm, rank test P = 0.008 and P = 0.452, respectively, Figure 5E). This suggests that mDia1 performs a function related to edge dynamics that is independent of F-actin assembly such as inhibiting barbed-end capping proteins (Shekhar et al., 2015) or regulating microtubules and cell-matrix adhesions (Ishizaki et al., 2001, Yamana et al., 2006).

In summary, these GC analyses unveil a dual role of mDia1 in distinct spatial locations, one as a direct F-actin assembly factor at the base of the lamellipodia, and another as an indirect modulator of edge motion through actin-independent functions.

### GC analysis detects causal shifts between wild-type and mutant of VASP deficient in actin assembly

Lamellipodial and lamellar F-actin assembly is regulated by numerous additional factors that are fine-tuning edge motion. In particular, the elongation factor VASP localizes to the tip of lamellipodia and accelerates elongation of actin filaments (Bear and Gertler, 2009). We took advantage of the vast biochemical knowledge of VASP-actin interactions to test whether our GC analysis has the sensitivity to detect subtle shifts in F-actin assembly and cell edge movement as a consequence of genetic mutations in VASP.

We co-imaged actin with SNAP-tagged wild-type VASP (VASP^WT^) and Halo-tagged S239D/T278E mutant VASP (VASP^MT^) in a VASP knock-down cell line (Figure S6A). VASP^MT^ has previously been described to maintain proper membrane localization, however with attenuated actin polymerization activity (Benz et al., 2009). In our U2OS cells, VASP^WT^ localized to a narrow band at the very tip of the lamellipodia (Figure 6A, Video S15). At large, this localization pattern applied also to VASP^MT^, but VASP^WT^ shifted towards the tip of the cell edge (Figure 6A). By CC analysis, we found that fluctuations in VASP^WT^ recruitment preceded edge protrusions by ∼3 s at the very front but were decoupled from edge motion in layers 2-4 (Figure 6B, only layers 1 and 2 are shown). The same spatiotemporal pattern arose for VASP^MT^ recruitment (Figure 6B), although the CC peak in layer 1 was reduced compared to VASP^WT^.

**Figure 6.**
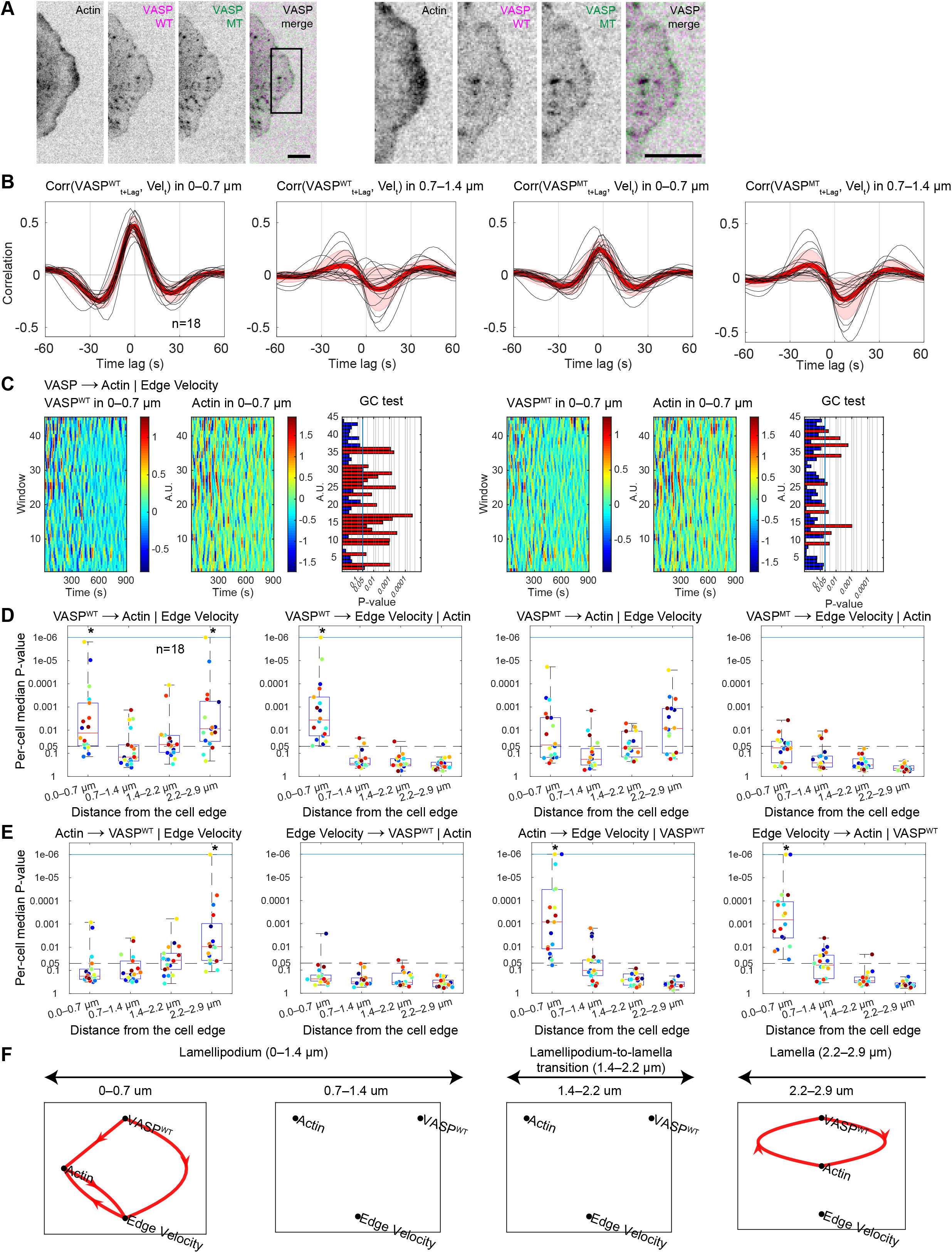
Granger-causality analysis of wild-type and mutant VASP co-imaged in the same cell. (A) Representative images of a U2OS cell (left) co-expressing mNeonGreen-actin, SNAP-tagged wild-type VASP (VASP^WT^) and Halo-tagged S239D/T278E VASP mutant (VASP^MT^). Scale bars, 5 µm. Right panel, enlarged insets. (B) Black curves, per-cell averaged cross-correlation of VASP^WT^ (left) and VASP^MT^ (right) with the edge velocity; red curves, population average (n = 18 cells); shaded bands, ±2 × SEM. (C) Activity maps of actin, VASP^WT^ (left) and VASP^MT^ (right), and associated P-values of GC from VASP^WT^ and VASP^MT^ to actin in the lamellipodia front for the cell in (A). Red, significant P-values. (D) Distributions of per-cell median P-values of Granger-causal relations for VASP^WT^ (left) and VASP^MT^ (right) (n = 18). The symbol (*), per-cell P-values significantly smaller than 0.05 (Wilcoxon signed rank test). (E) Granger-causal relations among VASP^WT^, actin and edge velocity as described in (D). (F) GC pathway diagrams between VASP^WT^, actin and edge velocity in the lamellipodia and lamella regions. See Figure S6 for GC pathway diagrams between VASP^MT^, actin and edge velocity. See also Video 15.

GC analysis identified marked differences between VASP^WT^ and VASP^MT^. Shown for a representative cell, VASP^WT^ recruitment G-causes F-actin assembly in most windows at the lamellipodia front (Figure 6C, median P-value 0.047) whereas VASP^MT^ recruitment was G-noncausal (median P-value 0.152). The differences were consistent over a cell population imaged over multiple independent experimental sessions (n = 18). VASP^WT^ was G-causal for F-actin assembly at the lamellipodia front (rank test P = 0.025), but VASP^MT^ was not (rank test P = 0.831, Figure 6D). This result shows the exquisite sensitivity of GC analysis in pinpointing differences between molecular activities that are not detectable by CC analysis.

Unexpectedly, GC analysis also identified a direct causal relation from VASP^WT^ to edge velocity, which was independent of the measured F-actin dynamics (rank test P < 0.001, Figure 6D). VASP^MT^ did not show this direct causal link (rank test P = 0.831, Figure 6D), indicating that the strong correlation between VASP^MT^ and edge velocity in Figure 6B does not imply causation.

Next, we used GC analysis to test possible feedback from F-actin to VASP recruitment (Figure 6E). Contrary to the feedback between F-actin and Arp2/3, which accompanies dendritic nucleation of actin filaments throughout the entire lamellipodia, VASP and actin are in bidirectional G-causal relations only in the most distal layer of the lamella (∼2.3–2.9 µm, Figure 6F). We suspect that this relation is a numerical artifact of the strong colocalization of VASP and F-actin at focal adhesions (Krause et al., 2003) (Video S15).

In summary, co-imaging of VASP^WT^ and VASP^MT^ first confirmed the biochemically characterized deficiency of VASP’s S239D/T278E mutation in F-actin assembly in a living cell and unveiled direct causal influence of VASP on edge motion. Notably, per our GC analysis, VASP-elongation of F-actin was confined to the lamellipodia front, without any feedback from F-actin assembly.

### Two discrete actin networks independently drive edge motion

Our GC pipeline delineated spatial zones in which Arp2/3, mDia1, and VASP assume differential G-causal roles in assembling F-actin. We wondered whether these three actin regulators operate in parallel or whether they are causally related among themselves. Specifically, we asked whether Arp2/3 and VASP operate cooperatively, i.e. VASP elongates F-actin nucleated by Arp2/3, or separately in differentially regulated F-actin networks (Winkelman et al., 2014, Pollard and Borisy, 2003). In the cooperative case, Arp2/3 recruitment would be expected as indispensable for explaining VASP recruitment (Figure 1E(3)). VASP-mediated F-actin assembly may also promote Arp2/3 recruitment (Figure 1E(4)), which would be reflected by a G-causal relation from VASP to Arp2/3. If Arp2/3 and VASP operated separately, our pipeline would report no G-causal relations.

To test these hypotheses, we co-imaged U2OS cells (n = 18) expressing SNAP-tagged exogenous VASP and endogenous Halo-tagged Arp3 (Figure 7A). For a representative cell, the GC tests at individual target windows at the lamellipodia front indicated that neither one of the two actin regulators are causal for the other (Figure 7B, median P-values > 0.222). The absence of such relations was further confirmed for lamellipodia and lamella at the level of the entire cell population (Figure S7A, rank test P > 0.204). On the other hand, VASP is identified as G-causal for edge motion, independent of Arp2/3 (Figure S7A, rank test P < 0.016), suggesting that Arp2/3 and VASP operate in two distinct F-actin architectures, which synergistically drive cell edge protrusion. In line with this prediction, GC pathway analysis indicated that Arp2/3 and VASP contribute as separate F-actin assembly factors to cell edge protrusion (Figure 7C). Importantly, this result does not invalidate the earlier established linear causal chain from Arp2/3 → actin → edge motion (Figure 2E). In the data underlying Figure 7C, actin is not observed and thus Arp2/3 appears causal for edge motion. Of note, the fluctuation signals of Arp2/3 and VASP correlated strongly with zero-time lag, which relates to their concurrent recruitment during P/R cycles (Figure S7B). However, co-recruitment does not mean coupled function. This underlines again the marked difference between correlation and causation analysis.

**Figure 7.**
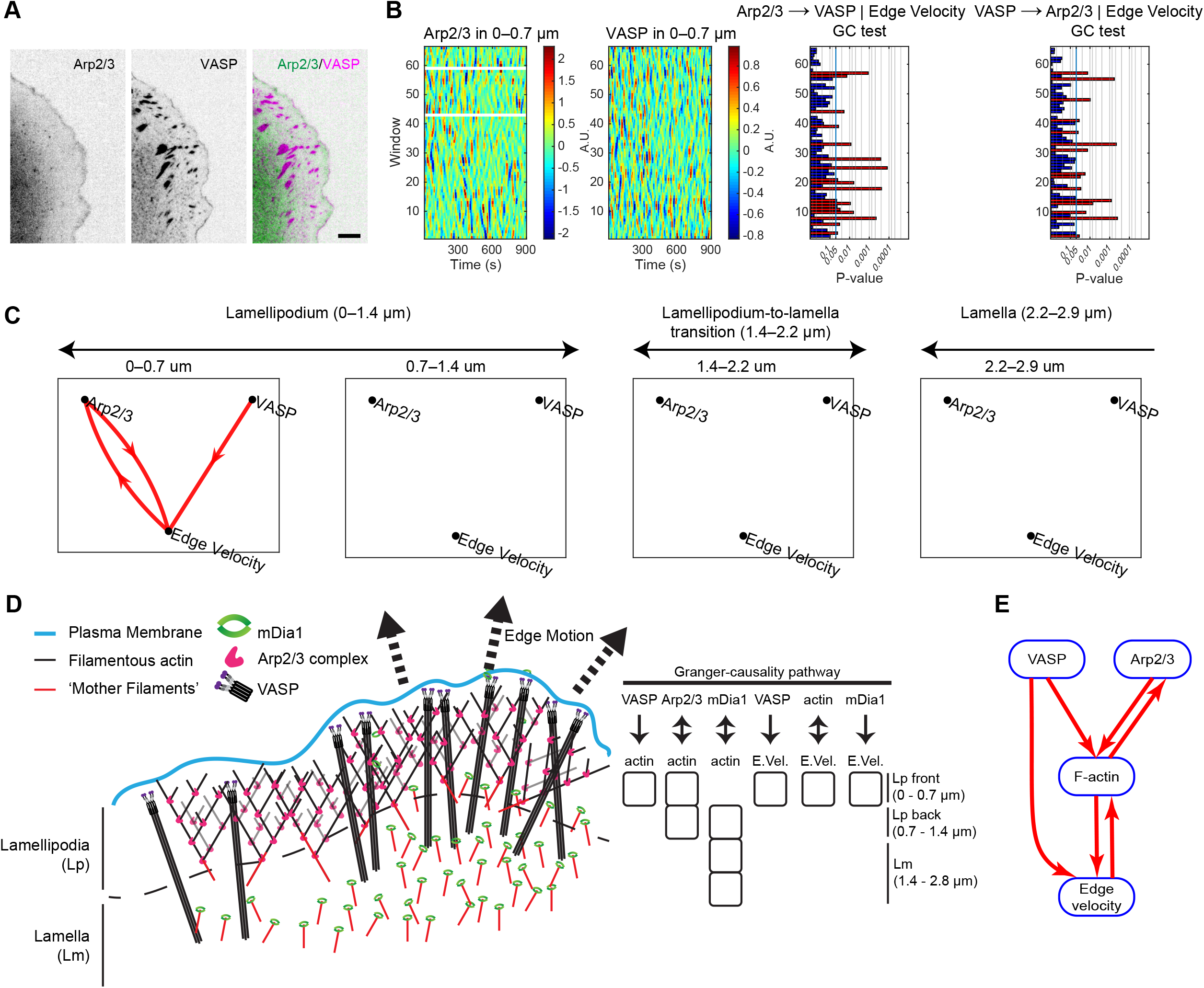
Two discrete F-actin networks drive edge motion. (A) Representative images of a U2OS cell co-expressing endogenously Halo-tagged Arp3 and exogenous SNAP-tagged VASP. Scale bars, 5 µm. (B) Arp2/3 (1^st^ panel) and VASP (2^nd^ panel) activities at the lamellipodia front of the cell shown in (A). P-values of GC from Arp2/3 to VASP (3^rd^ panel) and from VASP to Arp2/3 (4^th^ panel) in individual windows. Red, significant P-values. (C) GC pathway diagrams between Arp2/3, VASP and edge velocity in the lamellipodia and lamella regions. (D) Overview of G-causal relations of Arp2/3, mDia1, VASP and edge motion identified in the lamellipodia and lamella. Arp2/3 is Granger-causal for F-actin assembly only in the lamellipodia, VASP only in the lamellipodia front, and mDia1 in the lamellipodia back and lamella. No feedback is identified from F-actin assembly to VASP recruitment, while feedbacks are detected from F-actin to Arp2/3 and mDia1 in their respective activity areas. The GC analysis suggests that Arp2/3 and VASP operate independently to Granger-cause edge motion. We derive from this data the proposal that Arp2/3-mediated branched actin network and VASP-elongated linear actin bundles are independent F-actin network architectures synergistically driving edge motion. (E) Combined G-causal relations in the lamellipodia front between Arp2/3, VASP, actin and edge velocity.

## Discussion

When perturbed, regulatory pathways containing feedbacks and redundancies tend to respond with instantaneous adaptation. Thus, the ensuing phenotype is difficult to interpret in terms of the function the targeted component assumes in the unperturbed pathway. To reconstruct cause-effect relations between components of pathways, we adopt here the framework of Granger causality to live cell fluorescence imaging of the unperturbed system.

We chose the lamellipodial and lamellar actin assembly pathways to present three classes of experiments: i) control experiments with known outcome, validating the technology per se; ii) experiments with expected outcomes that were confirmed by GC analysis and thus raise confidence in the approach; iii) experiments that led to insights inaccessible by conventional perturb-and-observe approaches. Among the third class, our analyses define spatially segregated functions of mDia1 as an actin nucleator and as a nucleation-independent modulator of actin dynamics; and the existence of independently operating F-actin structures under the regime of Arp2/3 and VASP.

Our pipeline distinguishes with high specificity and sensitivity causation from correlation and allows the unmixing of functionally distinct molecular activities, which visually seem to be represented by identical fluorescence image fluctuations. This feature is illustrated with a VASP mutant that is deficient in actin-polymerization but localizes qualitatively identically to wildtype VASP at the lamellipodia edge. Our analysis recognizes that the VASP mutant is G-noncausal for F-actin assembly and edge motion, although its recruitment to the leading edge positively correlates with edge motion. We further identify that Arp2/3, actin and edge motion are in a linear causal chain, a finding in good agreement with the literature. Finally, our analyses determine that different actin regulators cause F-actin assembly in distinct spatial domains within the lamellipodia and lamella. Arp2/3 is shown to be G-causal for F-actin assembly only in the lamellipodia (0–1.4 µm from the edge), VASP only in the lamellipodia front (0–0.7 µm), and mDia1 at the lamellipodia base and in the lamella (0.7–2.9 µm, Figure 7D).

The Arp2/3 complex requires a pre-formed filament or “mother filament” to act as an actin branch nucleator (Mullins et al., 1998). The origin and source of the mother filament remains obscure. In two independent studies, the formin mDia1 was suggested to stimulate Arp2/3 activity *in vitro* and to precede lamellipodia protrusion onset *in vivo* (Isogai et al., 2015, Lee et al., 2015). Our data in U2OS cells determine that mDia1 initiates actin assembly during retraction and is G-causal for F-actin at the lamellipodia base and in the lamella (Figure 5F). Indeed, this may be the region where mother filament seeds form prior to protrusion onset, followed by autocatalytic nucleation of branched actin after activation and recruitment of Arp2/3 more proximal to the leading edge. In addition to mDia1’s selective G-causal relation for F-actin assembly at the lamellipodia-to-lamella transition, we found an unexpected direct G-causal relation between mDia1 and edge motion at the lamellipodia front, where this nucleator is G-noncausal for F-actin (Figure 5F). At first sight, this outcome could be interpreted as mDia1-decorated F-actin be too scarce compared to F-actin structures controlled by other nucleators for a causality between mDia1 and F-actin to be detected. However, our pipeline determined a causal link from mDia1 to edge motion not mediated through F-actin. Hence, the mDia1 signal does contain information to support the prediction of an actin-independent link to edge motion. If the mDia1 image fluctuations were purely noise, this link would not be extracted. We propose that the GC from mDia1 to edge movement relates to mDia1’s function in freeing F-actin barbed ends from capping proteins. This finding indicates the unique opportunities perturbation-free analyses generate to distinguish multi-functional properties of components. No perturbation experiment could be designed to determine this duality in mDia1’s action in the lamellipodia and lamella.

The proposed pipeline further identified a direct causal chain at the lamellipodia front from Arp2/3 to F-actin to edge velocity with feedback from F-actin to Arp2/3, and a chain from VASP to F-actin to edge velocity with a second G-causal link from VASP to edge velocity independent of F-actin-polymerization (Figure 7E).

We interpret this result with a model, in which the membrane-tethered VASP bundles F-actin (either by VASP alone or in cooperation with fascin) (Bachmann et al., 1999, Breitsprecher et al., 2008, Schirenbeck et al., 2006, Winkelman et al., 2014) and thus contributes to protrusion forces in parallel to VASP’s activity as an actin polymerase. Indeed, the direct link to edge motion is abrogated by VASP mutant that also abrogates F-actin bundling (Doppler and Storz, 2013) (Figures 6D and S6C). This model is also consistent with our finding that VASP- and Arp2/3-induced F-actin assembly G-causes edge motion independently from each other (Figure 7C) and highlights the sensitivity of our pipeline in deconvolving the inputs of distinct system components into a common target – information that would have been inaccessible using traditional perturbation approaches.

The performance of Granger causal inference is repeatedly disputed in the literature (Stokes and Purdon, 2017, Barnett et al., 2018, Stokes and Purdon, 2018). Where most discussions of these issues land is an agreement that the performance depends on the dataset and the underlying model of the connectivity of variables. Thus, it is imperative to validate the model assumptions. In most cases such a validation has to rely on empirical analyses, as the causal structure of the analyzed variables is generally unknown.

To accomplish this in our work, we then checked reproducibility of GC outcomes across independent imaging experiments with overlapping components: (actin, Arp2/3) vs (actin, cytoplasmic HaloTag) vs (actin, mDia1) vs (actin, VASP^WT^, VASP^MT^) vs (Arp2/3, VASP). The GC pathways identified from the partially observed systems were consistent enough to draw hypotheses for causal pathways in the whole system of Arp2/3, VASP, mDia1, actin and edge motion. To check how much the GC network outcome depends on the modeling strategy, we implemented variants of our framework in which one of the key strategies of our model was oppressed (see Methods and Figure S8). We lost consistency in the GC pathway outcomes across different imaging experiments when we eliminated spatial propagation of signals in the SPAR model or when we dropped a flexible regression model selection under the Akaike-Information Criterion (AIC) (see Methods). This shows that inclusion of the information from neighboring windows is critical to eliminate confounding factors, as is the proper choice locally of the time delays between variables. On the other hand, our tests indicated that the GC pathway prediction is insensitive towards transformation of the raw image signals into LFS signals (Figure S8).

The key limitation of the GC framework is that G-causal relations can only be determined within the system of co-observed variables (Eichler, 2013). While the proposed pipeline determines cause-and-effect relations between pairs of molecular processes, it is not yet able to map out directly causal interactions in larger process circuits with multiple components funneling information through one common component. Remedy to this limitation will arise from the development of hyperspectral imaging of an increasing number of components (Valm et al., 2017) and from expanded multivariate GC models that integrate data from several rounds of experiments under identical conditions but different configurations of component labeling.

## Supplemental information

Supplemental Figures S1–S8

Supplemental Videos S1–S15

Supplemental Tables S1–S2

## Acknowledgments

We thank Dr. Dick McIntosh (University of Colorado, Boulder, CO) for kindly providing the U2OS osteosarcoma cells, Dr. Tilmann Bürckstümmer and all Addgene depositors for sharing reagents and Dr. Dana Reed (UT Southwestern Medical Center) for her logistical support and laboratory management. We also thank the UT Southwestern BioHPC facility for providing high-performance computing systems. This study was supported by NIH grants K25EB028854 to JN and R35GM136428 to GD.

## Author contributions

Conceptualization: G.D; Methodology: J.N., T.I.; Formal analysis: J.N., T.I., Data Curation: J.N.; Validation: T.I., J.C., K.B.; Resources: T.I., J.C., K.B.; Investigation: G.D., T.I., J.N.; Supervision: G.D.; Funding acquisition: J.N., G.D.; Writing – original draft: J.N., T.I., G.D.; Writing – review & editing: J.N., T.I., K.B., G.D.

## Declaration of interests

The authors declare no competing interests.

## Star Methods

### KEY RESOURCES TABLE

| REAGENT or RESOURCE | SOURCE | IDENTIFIER |
| --- | --- | --- |
| **Antibodies** |  |  |
| Mouse anti-β-actin | Sigma | AC-15; A1978 |
| Mouse anti-mDia1 | BD Biosciences | 610848 |
| Rabbit anti-VASP | Cell Signaling | 3112 |
| Mouse anti-ARP3 | Santa Cruz Biotechnology | B-1; sc-374200 |
| Rabbit anti-HaloTag | Promega | Rabbit: G9281 |
| Rabbit anti-SNAP | New England BioLabs | P9310S |
| **Chemicals, peptides, and recombinant proteins** |  |  |
| DMEM | Gibco | 12430054 |
| Trypsin/EDTA | Gibco | 15400054 |
| DMEM, high glucose, HEPES, no phenol red | Gibco | 21063029 |
| Bovine Serum Albumin | Equitech-Bio | 190524 |
| Fibronectin | Sigma | F1141 |
| Polyethylenimine (PEI) | Sigma | 408727P |
| Puromycin | Gibco | A1113802 |
| HaloTag TMR ligand | Promega | G8251 |
| SNAP-Cell 647-SiR | New England BioLabs | S9102S |
| Janelia Fluor 549 (JF549) HaloTag ligand | Promega | GA1110 |
| **Experimental models: cell lines** |  |  |
| Human Osteosarcoma U2OS cells | Laboratory of R. McIntosh,<br>University of Colorado<br>Boulder, CO | N/A |
| **Oligonucleotides** |  |  |
| See Table S1 | N/A | N/A |
| **Recombinant DNA** |  |  |
| pSpCas9(BB)-2A-GFP (PX458) | (Ran et al., 2013) | Addgene 48138 |
| pSpCas9n(BB)-2A-Puro (PX462) | (Ran et al., 2013) | Addgene 48141 |
| pX462-ACTR3-FR1 (ARP3) | This paper | N/A |
| pX462-ACTR3-FR2 (ARP3) | This paper | N/A |
| pX458-DIAPH1 (mDia1) | This paper | N/A |
| pMA-tial1 | (Lackner et al., 2015) | N/A |
| pMA-tial1-HaloTag-N-18-ACTR3 donor | This paper | N/A |
| pMA-tial1-mNG2<sub>11</sub><sup>4</sup>-C-18-DIAPH1 donor | This paper | N/A |
| psPAX2 | Trono Lab Packaging and<br>Envelope Plasmids<br>(unpublished) | Addgene 12260 |
| pMD2.G | Trono Lab Packaging and<br>Envelope Plasmids<br>(unpublished) | Addgene 12259 |
| pLVXCMV100 | (Dean et al., 2016) | Addgene 110718 |
| pLVXCMV100-mNeonGreen-18-actin | This paper | N/A |
| pLVXCMV100-SNAP-21-VASP | This paper | N/A |
| pLVXCMV100-Halo-21-S235D/T273E | This paper | N/A |
| pLVXCMV100-SNAP-21-mDia1 | This paper | N/A |
| pLKO.1 – TRC cloning vector | Moffat et al., 2006 | Addgene 10878 |
| pLKO.1-shVASP TRCN0000369620 | This paper | N/A |
| pLVX-IRES-Puro | Clontech | 632183 |
| pLVX-IRES-Puro-mNeonGreen2<sub>1-10</sub> | This paper | N/A |
| **Software and algorithms** |  |  |
| Matlab | The MathWorks, Inc. | https://www.mathworks.com/products/matlab.html |
| Windowing-Protrusion Package in Matlab | (Ma et al., 2018) | https://github.com/DanuserLab/Windowing-Protrusion |
| Granger-causal pathway inference of lamellipodium regulatory system | This paper | Zenodo:<br>https://doi.org/10.5281/zenodo.6374351 |
| CRISPOR | (Concordet and Haeussler, 2018) | http://crispor.tefor.net/ |
| Fiji | (Schindelin et al., 2012) | https://fiji.sc/ |

### RESOURCE AVAILABILITY

#### Lead contact

Further information and requests for resources and reagents should be directed to and will be fulfilled by the Lead Contact, Gaudenz Danuser.

#### Materials availability

Requests for reagents will be fulfilled by contacting the Lead Contact upon reasonable requests.

#### Data and code availability

- Microscopy data reported in this paper will be shared by the lead contact upon reasonable request.
- All original code including a user-friendly Graphical User Interface has been deposited at Zenodo and GitHub, and is publicly available as of the date of publication (https://github.com/DanuserLab/Granger-Causality-Analysis-of-Lamellipodia). DOIs are listed in the key resources table.
- Any additional information required to reanalyze the data reported in this paper is available from the lead contact upon request.

### EXPERIMENTAL MODEL AND SUBJECT DETAILS

#### Cell lines

Human Osteosarcoma U2OS cells were a kind gift from Dr. Dick McIntosh (University of Colorado Boulder) and were maintained in DMEM supplemented with 10% fetal bovine serum (Sigma; F0926-500ML) in a humidified incubator at 37 °C and 5% CO2. All cells were tested for mycoplasma using a PCR-based Genlantis Mycoscope Detection Kit (MY01100). Cells were not authenticated.

### METHOD DETAILS

#### Plasmids

pSpCas9n(BB)-2A-Puro (PX462), pSpCas9(BB)-2A-GFP (PX458) and pLKO.1 were from Drs. Feng Zhang and David Root, respectively (Addgene plasmids #48141, #48138 and #10878). Gene-targeting single guide RNAs (sgRNAs) were designed using CRISPor (Concordet and Haeussler, 2018). The self-cleaving donor vector pMA-tial1 was a kind gift from Dr. Tilmann Bürckstümmer (Lackner et al., 2015). HaloTag flanked with homologous arms targeting the last exon of Arp3/ACTR3 (NM_005721) was cloned seamlessly into pMA-tial1 (HaloTag-N-18-ACTR3 donor) using HiFi Assembly (NEB). A codon-optimized split fluorescent protein mNeonGreen2_1-10_ (mNG2_1-10_) was cloned with SalI/NotI into the XhoI/NotI sites of pLVX-IRES-Puro (Clontech). mNeonGreen2_11_ placed in tandem of four (mNG2_11_^4^), each separated with a GSGSG 5 amino acid linker and flanked with homologous arms targeting the first exon of mDia1/DIAPH1 (NM_00521), was blunt cloned into the EcoRV site of pMA-tial1 (mNG2_11_^4^-C-18-DIAPH1 donor). mNeonGreen, SNAP, Halo, human β-actin, mouse VASP and mouse mDia1 DNA fragments were PCR amplified with corresponding flanking homology regions to seamlessly clone into pLVXCMV100 using HiFi Assembly to generate mNeonGreen-18-actin, SNAP-21- and Halo-21-tagged VASP, and SNAP-21-mDia1 where 18 and 21 denotes the number of amino acids in the linker between the tag and the tagged gene. These linkers and the terminal ends used for tagging were chosen based on our previous work (Lee et al., 2015). The 18-linker sequence was (SGLRSGSGGGSASGGSGS) and the 21-linker sequence was (EPTTEDLYFQSDNAIAGRPRSSG). The S235 and T273 residues (corresponding to the S239/T278 residues on human VASP) in the Halo-tagged VASP were PCR mutated to aspartic acid and glutamic acid, respectively. VASP knockdown (KD) cells were generated and enriched by lentiviral infection of pLKO.1 containing the short-hairpin RNA targeting VASP (TRCN0000369620) and by subsequent puromycin (2 µg/ml) selection, 48 hours post-infection. All new constructs used in this study were sequenced verified and are listed in Key Resources Table. Primers and gBlocks used for cloning are listed in Supplementary Table S1. mNeonGreen is licensed by Biotechnology & Pharmaceuticals, Inc.

#### Cell Culture

Cells were counted using Cellometer Auto 1000 Bright Field Cell Counter (Nexcelom). Lentiviral particles were generated using the packaging vectors psPAX2 and pMD2.G (Addgene plasmids #12260 and #12259). Infected cells were bulk sorted using FACS, or selected with Puromycin (1 µg/ml; Gibco).

#### Engineering of Genome-edited Cells

Arp3 was C-terminally tagged with HaloTag at its endogenous genetic loci in U2OS cells, and were generated as follows: 1 x 10^5^ U2OS cells seeded in a 6 well plate were transiently transfected overnight with two pX462 plasmids containing a ACTR3-targeting sgRNA together with HaloTag-N-18-ACTR3 donor using polyethylenimine (PEI; Sigma). Genome-edited cells were labeled with Halo-TMR ligand (5 µM; NEB), and single cell Fluorescence Activated Cell Sorted (FACSed) into 96 wells tissue culture plates. Genome-edited cells expressing Halo-Arp3 were screened and verified with Western Blotting.

Initially, we endogenously tagged mDia1 with a single HaloTag, but the signal from the tagged protein was too dim, presumably due to the low abundance of mDia1 molecules in cells. This resulted in a very low SNR not suitable for imaging and causality inference (data not shown). To overcome this limitation, we used a self-complementing split fluorescent protein mNeonGreen2_1-10/11_ system with reduced background (Feng et al., 2017), which allows signal amplification of low abundant proteins using a relatively short, 16 amino acid-long 11th β-strand. mDia1 was N-terminally tagged with four mNG2_11_ peptides placed in tandem (mNG2_11_^4^), each separated with a 5 amino acid linker (GSGSG), totaling 79 amino acids in size from its endogenous loci in cells expressing the mNG2_1-10_ cells: 1 x 10^5^ U2OS-mNG2_1-10_ cells seeded in a 6 well plate and transiently transfected overnight with plasmid pX458 containing a DIAPH1-targeting sgRNA together with the mNG2_11_4-C-18-DIAPH1 donor vector using PEI. Genome-edited cells were bulk FACSed, containing both heterozygous and homozygous edited cells, verified with Western Blotting and Immunofluorescence using anti-mDia1 antibodies (Figures 5A and S5).

#### Introduction of Fluorescent Probes and Considerations to Minimally Perturbing the Cells

To minimize potential unforeseen perturbations, both ACTR3 and DIAPH1 CRISPR donor vectors contain an additional 18 amino acid linker (SGLRSGSGGGSASGGSGS) between the tag and the tagged gene inspired by Dr. Michael Davidson’s designs. Lentiviral constructs harboring a previously described truncated CMV promoter containing only the first 100 bps of the promoter (CMV100) (Dean et al., 2016) were used to achieve low expression of the exogenous genes. In addition, all the tagged genes contained a variable amino acid linker varying from 18-21 amino acids to minimize interference due to tagging.

#### Live-Cell Time-lapse Imaging of Cells

Cells were seeded on fibronectin (10 µg/ml)-coated #1.5 glass-bottom dishes and allowed to spread overnight. The following day, cells were labeled with HaloTag ligand conjugated with JF549 (0.4 – 1 µM; Promega) and/or SNAP ligand conjugated with SiR-647 (0.25 – 1 µM; NEB) for 30 minutes, and subsequently washed twice with DMEM. Prior to imaging, the media was replaced with phenol-red free DMEM supplemented with 20 mM HEPES pH7.4. Time lapse image sequences were acquired on a climate-controlled (maintained at 37°C), fully motorized Nikon Ti-Eclipse inverted microscope with Perfect Focus System, equipped with a 60×, 1.49 NA APO TIRF objective (Nikon) with an additional 1.8× tube lens (yielding a final magnification of 108×; Andor Technology), and an Andor Diskovery illuminator coupled to a Yokogawa CSU-X1 confocal spinning disk head with 100 nm pinholes. Image sequences were recorded using a scientific CMOS camera with 6.5-µm pixel size (pco.edge) at a 3 Hz frame rate.

#### Overview of the Granger Causality (GC) analysis pipeline

The input to the Granger Causality (GC) pipeline is a set of two-channel fluorescence live cell movies visualizing the dynamic recruitment of proteins implicated in regulating cell edge protrusion and retraction cycles. Conceptually, GC analysis is not limited to protein recruitment, but any variable for which a time series can be extracted from a live cell movie can be processed, as discussed in the following sections. Given the specific test application in this work, one variable unrelated to protein recruitment is the local edge movement. We thus refer to the variables more generally as molecular and cellular activities. After image processing and statistical causal inference, the pipeline produces diagrams of the cause-effect relation network between the imaged activities. The pipeline is designed to automatically optimize computational parameters in a data-driven fashion. Thus, the pipeline runs from the raw movies to the cause-effect network with very few user inputs. In the following, we describe each step of the pipeline with the necessary detail.

In comparison to our previous analyses of edge motion and molecular activities (Ma et al., 2018, Azoitei et al., 2019), we upgraded the image and signal processing to obtain more accurate subcellular readouts, which was essential to the application of statistical modeling and causal inference. Specifically, we developed a new segmentation method called Multi-Scale-Automatic (MSA) segmentation that combines multiple segmentation results from different algorithms and parameters to generate accurate cell masks. In terms of signal processing, we introduced a decomposition of the molecular activity oscillation into low-frequency and high-frequency activities, because the molecular recruitments encompassed a sub-spectrum of oscillations that were much slower than those of the protrusion-retraction (P/R) cycles. These technical improvements contributed to generating robust and consistent GC predictions within the network of actin regulators.

#### Cropping an active part of a cell showing dynamic edge motion

Several cells were typically imaged in one fluorescence movie. For the statistical inference of causal relations among molecular activities near the edge, we cropped a part of a cell where most of the cell edge segments displayed at least a few cycles of P/R events. In such an area, the imaged proteins typically showed multiple cycles of recruitments co-fluctuating with the dynamic edge motion. We avoided cropping regions with frequent occurrence of ruffling, because the fluorescence intensities at ruffles are abnormally amplified. We also avoided regions that were filled with static cytoskeletal structures such as transversal arcs in the actin images and focal adhesions in the VASP images. Unlike protein recruitment in the lamellipodia, the molecular activities on the static structures were unrelated to edge movement. These cropping criteria were equally applied across all the fluorescence movies of different proteins.

#### Multi-Scale-Automatic (MSA) segmentation

The cell area in each frame of cropped live cell movies was segmented to extract cell edge motion profiles and molecular activities near the cell edge. The previous segmentation methods developed in our lab for edge motion profiling consist of three steps: (i) smoothing images with a Gaussian filter; (ii) using an automatic intensity thresholding algorithm; (iii) morphological closing of cell masks for filling small holes. The automatic thresholding algorithms include Otsu (Otsu, 1979), Rosin (Rosin, 2001), and a custom-written algorithm called ‘minmax’ as previously described (Ma et al., 2018). These methods were successfully applied to a variety of imaging studies with different data quality (Machacek et al., 2009, Han et al., 2015, Ma et al., 2018, Azoitei et al., 2019). However, the automatic thresholding algorithms often failed when the signal-to-noise-ratio varied between different parts of the images, and it was inconvenient for a user to have to optimize the segmentation parameters for each movie.

To automate the manual optimization of the segmentation parameters and to improve the accuracy of edge motion profiles, we developed the MSA segmentation method, which is a simple ensemble-learning method (Rokach, 2010) that combines the previous segmentation methods in our lab. First, we computed 42 different cell masks for each image by applying all combinations of segmentation parameters: 3 thresholding algorithms, 7 different σ values of Gaussian filters for smoothing ranging from 0 to 3 pixels, and 2 different radius parameters for morphological closing. For each pixel, we then computed the mask score which is defined by the number of different cell masks that classified the pixel as foreground. The mask scores themselves across pixels became a new, transformed image where the signals are proportional to foreground probabilities. The mask score image was much easier to segment because the scores in the background clustered at a low value and the scores in the foreground clustered at a high value. We used the middle value of the low and high scores of the two peaks to segment the mask score images. In most cases, the MSA segmentation generated more accurate cell masks than any single combination of the thresholding algorithms and Gaussian filtering parameters, as assessed by visual inspection.

To segment the two-channel fluorescence images with different combinations of proteins as objectively as possible, we added the two images of the different proteins so that the cell masks did not depend on a particular protein. Since our two-channel images had pixel intensities in a similar range, the addition was applied without scale adjustment. The summation image was segmented using the MSA segmentation method.

#### Cell edge trembling correction

To ensure accurate edge motion profiling, we investigated whether cell masks of a movie measured the morphological changes in an unbiased manner. We occasionally found events, in which cell masks globally dilated or eroded by ∼1 pixel between two consecutive frames before stabilizing again at the following frames. This cell edge trembling was subtle but caused a systematic bias in the cross-correlation between edge velocity and molecular recruitment at lag 0, as previously addressed (Marston et al., 2020). Though the MSA segmentation reduced this edge trembling effect, the bias in the cross-correlations remained.

We corrected the segmented edge trembling effect by introducing weighted averaging of three consecutive cell masks. In particular, we first computed the level set function at each location (*x*, *y*) defining the distance from the cell boundary toward the cell inside. The level set functions, *ϕ*_*t*_(*x*, *y*) at three consecutive time points were averaged with weights as follows:

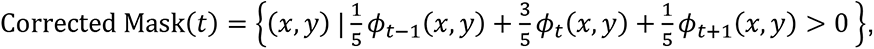

where the weights were chosen heuristically. The corrected masks generated subcellular edge velocities and molecular activities that showed no bias in the cross-correlations.

#### Tracking edge motion and probing windows

The segmented cell boundaries of a live cell movie allowed us to measure edge velocities and molecular activities at a submicron-scale using the previously developed morphodynamic profiling algorithms (Ma et al., 2018, Machacek et al., 2009). We tracked the segmented cell boundaries and small probing windows that partitioned the area near the edge (Figure 1G). In the application of the algorithm, we used the option ‘Constant number’ as a method of propagating the windows between consecutive time points. To select the size of the probing windows, we first visually inspected an overall depth of the lamellipodia from the edge, which was ∼1.4 µm for most of the imaged cells. We decided to analyze a band 0–2.9 µm sub-adjacent to the cell edge in order to capture probable differences in molecular recruitment between lamellipodia and lamella. Lamellipodia and lamella were further split in two layers in order to capture potential gradients in regulation. This determined the size of the probing windows to 720×720 nm^2^ width and depth (Figure 1G). Using this window grid, we extracted time series of local edge velocity and recruitment of two different proteins.

#### Subtraction of low-frequency oscillation for molecular recruitments

To test the requirement of GC-based causal inference and correlation analysis for compatible time scales between the analyzed variables, we determined the characteristic time scales of extracted edge velocities and molecular recruitments using auto-correlation analysis. In each probing window, the auto-correlation function (ACF) of an edge velocity time series was computed. The ACFs were averaged over the windows for each cell. The per-cell averaged ACFs were remarkably consistent (Figure 3A), showing that the average duration of P/R cycles is maintained across experiments. The ACFs had a maximal negative value at ∼30 s on average (Figure 3A), indicating a duration of P/R cycles of ∼60s. In contrast, the ACFs of molecular recruitments indicated much slower cycles with durations > 240 s (Figure 3A for actin and Arp2/3).

We decided to cancel out fluctuation components much slower than the average P/R cycle, assuming that these components are marginally involved the regulation of edge movements. To accomplish this we computed moving medians over rolling time intervals of 60 s (20 frames), i.e. the average duration of a P/R cycle. Examples of raw molecular activity time series (*X*_*t*_) and their low-frequency components (*LF*_*t*_) are shown in Figures 3B and 3D. Taking *LF*_*t*_ as baseline activity, the low-frequency subtracted (LFS) molecular activity was defined as the percentage of activity above baseline relative to the baseline:

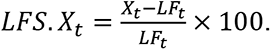

Intriguingly, the LFS activities of actin and Arp2/3 recruitments displayed spatiotemporal dynamics similar to the corresponding edge velocity dynamics (Figures 3C and 3E). We concluded that the LFS activities are the primary drivers of the edge fluctuations and thus focused our causality analysis of actin regulating factors on this signal.

#### Reconstruction of synthetic live cell movies using LFS-molecular recruitments

To examine the difference between molecular recruitments observed in the fluorescence movies and computed LFS-molecular activities, we reconstructed synthetic live cell movies that visualized the LFS-activities next to the original fluorescence intensities (Video S4).

The process of mapping LFS time series back into the image space required a few steps to ensure the smooth transition between the window-based array of LFS-activities and time-lapse images. Since the LFS-activities had signed values, they were rescaled to the [0, 1] range. We matched the 1^st^ percentile of the activities to 0 and the 99^th^ percentile to 1, and truncated at the two extreme percentiles to exclude outliers. Since a cell shape could not be perfectly morphed into the shape at the next time frame, the trajectory of locations of each window over time was not smooth enough, so direct mapping of the scaled LFS-activities into the pixels covered by the associated probing window in in the image space generated artifacts. Instead, we applied cubic smoothing spline interpolation to the mapped pixel intensities using the Matlab function *csaps()* with a strong smoothing parameter. This resulted in successful reconstruction of LFS-activity movies from the original imaging data.

The reconstructed live cell fluorescence movies allowed us to visualize the activities of actin and its regulators that fluctuated at the time scale of edge P/R cycles. The movies showed a much stronger coupling of, for example, LFS-actin activity to P/R events than the original actin activities (Video S4).

#### Filtering out probing windows at quiescent edges

Before inferring GC relations, our pipeline implemented diagnostic tests that checked whether the imaged cell edge motion displayed systematic P/R events and whether the edge velocity time series were appropriate input for statistical time series analysis. Because our study targets pathways that regulate P/R cycles in migrating cells, we eliminated windows that sampled molecular activities in quiescent edge sectors. To identify such windows, we statistically tested if the edge velocity time series represented a white-noise process with the underlying assumption that actively regulated time series cannot be described by white noise. We used the Matlab function *lbqtest()* that implements the Ljung-Box test (Ljung and Box, 1978) to determine whether the auto-correlation of the edge velocity time series were zero – an indication that the edge in that sector was quiescent. If the proportion of the quiescent windows was > 50%, then the cell was labeled as overall quiescent and excluded from the subsequent GC inference.

For non-quiescent windows we also checked if the edge motion fulfilled stationarity. Since the GC analysis requires that the observed time series is free of a trend, we performed for each edge velocity time series an augmented Dickey–Fuller test (Dickey and Fuller, 1979) (using the Matlab function *adftest()*) – the standard test to identify the existence of systematic trends in time series data). If the majority of probing windows was determined to possess systematic trends, then the cell edge motion was determined as non-stationary overall and the data was excluded from further analysis.

#### Spatially propagating auto-regressive (SPAR) models for one-channel live cell movies

Inferring the GC relations within a dynamic system relies on a model that specifies how the time series data in the system are generated. To build such a model for live cell imaging data, we began with modeling the dynamics of molecular activities of one protein across subcellular windows and over time. On top of a purely auto-regressive model quantifying the dependency of a variable’s future on its past, we added spatial dependency to quantify how much the future of a variable in one location relates to the present and past of the variable in the four adjacent probing windows.

We denote by 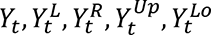 the molecular activities at time *t* in a target window and in its left, right, upper and lower neighboring windows, respectively. Our generative stochastic model for one protein, called the Spatially Propagating Auto-Regressive (SPAR) model, is written as follows.

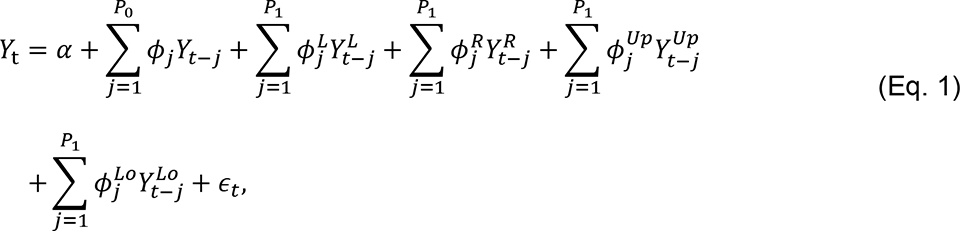

where *ε*_*t*_’s are independent identically distributed random errors with mean 0 and variance *σ*^2^, *P*_0_ is the auto-regressive order for the same window, *P*_1_ is the cross-regressive order for the four adjacent windows, and 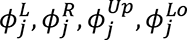 are the coefficients to quantify the influence of the left, right, upper and lower window time series, respectively, on the target series.

The number of possible SPAR models depends on the range of values that must be considered as plausible regressive orders, *P*_0_ and *P*_1_. For the present study, we considered *P*_0_ and *P*_1_ up to half the duration of an average P/R cycles, which is ∼30 sec or 10 time frames. Thus, we set 1 ≤ *P*_0_, *P*_1_ ≤ 10. This led to 10 × 10 possible SPAR models to describe the dynamics of one variable. As discussed next, these models are evaluated independently in every window, which comes with substantial computational burden. Therefore, to reduce the number of possible model orders for efficiency, we made the assumption that the cross-regressive order is less than the autoregressive order for the same window (*P*_0_ ≥ *P*_1_), because 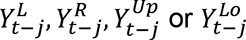 would be less correlated with *Y*_*t*_ than *Y*_*t*−*j*_ would be. This assumption reduced the number of candidate models from 100 to 55. After the model selection procedures described below, the selected orders ranged from 4 to 9 for *P*_0_, and from 1 to 3 for *P*_1_, for most molecular activities analyzed in this paper.

Our pipeline includes an exhaustive model selection step, which is essential for accurate causal inference (see below). In each probing window, we fitted all candidate SPAR models across the set range of regressive orders. The model parameters were estimated by the standard least-squares method based on which we computed the Akaike-Information Criterion (AIC) (Akaike, 1974) for all the fitted models. Analysis of the spatial distribution of the orders indicated that while the AIC values varied between cells and between activity, they were similar for the probing windows in one layer. Thus, we selected the optimal *P*_0_ and *P*_1_ as those minimizing the averaged AIC values per window layer and per-cell. This simplification is also necessary to limit computational cost and to increase robustness in model selection as not every window offers sufficient signal to noise ratio for a complete optimization of the AIC.

After AIC-based model selection, we statistically tested whether fluctuations in a particular molecular activity propagated between adjacent windows. For example, we found that fluctuations in actin recruitment near the cell edge actively propagated to adjacent windows while fluctuations in the concentration of a cytoplasmic probe showed no intracellular information flow (Figures 4A and 4B). To identify these information flows, we compared the prediction capability of the full model (Eq. 1) with the capability of reduced models in which the influence of activities in either the left, the right, the upper, or the lower window adjacent to the target window was excluded (Figure S1C). For example, to identify the information flow from the left window to the target window, we tested the following null hypothesis for the SPAR model in (Eq. 1):

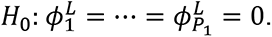

As discussed below (Eq. 3), this hypothesis is accepted, i.e. no information flow, or rejected, i.e. information flows from the left window to the target window using a Fisher test that compares the residuals of the full and reduced models. We obtained the P-values for four directional intracellular information flows at individual windows. These multiple testing results were then adjusted to control the false discovery rate (FDR) (Benjamini and Hochberg, 1995) at 20%.

#### Granger-causality test at a single window for two-channel live cell movies

We extended the SPAR model for the dynamics of one protein (Eq. 1) to identify inter-molecular information flows at each probing window. In the SPAR model for two-channel live cell movies, we specified the spatiotemporal dependency between subcellular molecular activities and local edge velocities as follows:

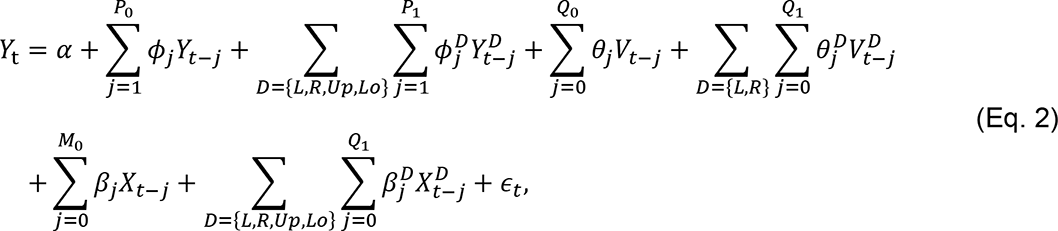

where *Y*_*t*_, *X*_*t*_ denote the putative effector and causative activities at time *t* (*t* = 1, 2, …, *T*), *D* = {*L*, *R*, *Up*, *Lo*} denote the left, right, upper, and lower windows to the analyzed target window, *V*_*t*_ denotes the edge velocities at the edge segment closest to the target window, *P*_0_, *P*_1_, *Q*_0_, *Q*_1_, *M*_0_ are the temporal lag orders for different features, and *ε*_*t*_’s are independent random errors with mean 0 and variance *σ*^2^ (Figure 2B).

In this SPAR model, a current activity of the effector (*Y*_*t*_) is explained by all the local features available in the two-channel movies. Importantly, in contrast to typical time series regression models, we include the activity at present time of the putative cause (*X*_*t*_) and thus refer to our analysis as *instantaneous* Granger-causality (Runge, 2018). Given that our imaging frequency is set to one multi-channel acquisition every 3 seconds, the activities of the putative cause and relevant features at the present time 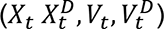 in (Eq. 2) can be regarded as the proxy of their past values up to ∼1.5 second before the processed time *t*. Incorporating a contemporaneous link turned out to be critical for the model to result in GC outcomes that are consistent with the known biochemical relations between Arp2/3 and F-actin. This indicates that inter-molecular information flows occur faster than multi-channel light microscopy permits without introducing photo-toxicity and bleaching.

Similar to the SPAR model for one protein, the optimal model orders were selected using the AIC. Orders up to the half of the P/R cycle were considered under the following constraints: 1 ≤ *P*_1_ ≤ *P*_0_ ≤ 10, 0 ≤ *Q*_1_ ≤ *Q*_0_ ≤ 10, 0 ≤ *M*_0_ ≤ 10. Since an exhaustive testing of all combinations of possible orders is computationally too costly, we employed a sequential model selection approach, where (i) the SPAR model in (Eq. 1) was first considered to select the auto-regressive orders, *P*_0_ and *P*_1_; (ii) using the selected *P*_0_ and *P*_1_, the model in (Eq. 2) without the terms of {*X*_*t*_} was used to choose the optimal *Q*_0_ and *Q*_1_; (iii) lastly, the optimal *M*_0_ was selected using the other pre-selected orders. As in the model selection for one protein, the model orders were chosen to be the one minimizing the averaged AIC values per probing window layer on a cell. In the example of SPAR models testing the GC from Arp2/3 to F-actin in the region 0–0.7 µm, the selected model orders varied among the analyzed 20 cells within ranges 6 ≤ *P*_0_ ≤ 8, 1 ≤ *P*_1_ ≤ 3, 1 ≤ *Q*_0_, *Q*_1_ ≤ 2, 0 ≤ *M*_0_ ≤ 3. It is noteworthy that the optimal lag orders between putative cause and effect relations were mostly *M*_0_ = 1 and models with only a contemporaneous term (*M*_0_ = 0) occurred for two out of 20 cells.

To identify the GC relations between two molecular processes in a particular window, we tested whether the activities of the putative cause in the same window ({*X*_*t*_}) were indispensable in explaining the current effector activity (*Y*_*t*_). In other words, we tested the null hypothesis for the SPAR model in (Eq. 2),

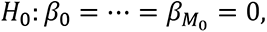

by using the following F-test statistic,

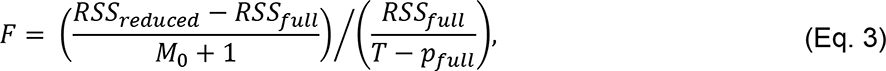

where *RSS*_*full*_ (*RSS*_*reduced*_) is the residual sum of squares of the full model (the reduced model or the model restricted by the null hypothesis *H*_0_), *T* is the total number of time points, and *p*_*full*_ is the number of parameters of the full model. A large F-statistic indicated the indispensability of the putative cause at a probing window, resulting in a small P-value. The P-values obtained by the F-tests in individual windows provided subcellular evidence for the tested causal relation.

The directionality of the GC relation follows from the temporal structure designed in the model for inference. Since we employ an instantaneous GC formalism the interpretation of directionality needs attention in the situation, in which only the contemporaneous link from *X*_*t*_ and *Y*_*t*_ is significant without any lagged dependency between (*X*_*t*−*j*_, *Y*_*t*_) or (*Y*_*t*−*j*_, *X*_*t*_), *j* ≥ 1. In this situation, the framework may be detecting bi-directional GC relations between {*X*_*t*_} and {*Y*_*t*_}, that is, a causal feedback loop. However, since the sampling rate permitted by microscopy may not always be fast enough, a significant contemporaneous link without support from a lagged dependency is insufficient to claim causal feedback. This situation never occurred in the presented image data.

### Statistical diagnostics of the Granger-causality tests in individual windows

After fitting the regression models and implementing the F-tests, we applied three diagnostic measures to validate the model estimates in individual windows. First, we checked auto-correlations of the residuals, which need to be close to zero at all lags. Second, we checked the condition number which is an index of multi-collinearity of the predictors in the regression model (Eq. 2). A large condition number (conventionally > 30) indicates severe multi-collinearity and unstable parameter estimation. Third, we checked whether the distribution of the residuals follows a normal distribution using the Kolmogorov-Smirnov (KS) test via the Matlab function *kstest()*. If those diagnostic measures indicate poor regression estimates, then the corresponding F-test P-values were not accepted for further cell-based analysis.

#### Granger-causal relations based on cell-to-cell variability

Following GC-analysis in each probing window, the per-window GC P-values were then integrated into per-window layer/per-cell median P-values. Because the elementary unit of our imaging experiments was one cell, the per-cell measurements were statistically tested to determine the final conclusion on GC relations based on cell-to-cell variability. We utilized Wilcoxon signed rank tests to determine whether the per-cell median P-values of *n* cells were significantly smaller than the nominal level 0.05, or equivalently, whether the majority of probing windows in a particular layer showed significant evidence for a putative causal relation over multiple independent cells. Using the Matlab function *signrank()*, the signed rank tests were implemented separately to the four layers that partitioned the lamellipodia and lamella region according to the distance from the cell edge (Figure 2D). The obtained rank test P-values in this study were listed in Supplementary Table S2.

#### Drawing GC pathway network diagrams

The GC analysis of two-channel live cell movies determined the combinatorial GC relations between the recruitment of two molecules and the edge motion dynamics at different distances from the edge (Figure 2D). The obtained GC relations resulted in connectivity networks, which we visualized using the hierarchical graph drawing algorithm (Sugiyama et al., 1981), which is implemented in the Matlab function *digraph()*. For all graphs we set the option of layout to *‘layered’*, assuming that edge motion is the ultimate effector of the pathways. It automatically generated GC networks at the different distances from the cell edge (Figure 2E), which unveiled spatially heterogeneous actin regulatory pathways in lamellipodia and lamella.

### Key modeling strategies which enable reproducible GC pathway outcomes

We chose to examine how much the GC outcomes varies between different modeling strategies. We implemented variants of our framework by removing one of the key modeling strategies at the time. We eliminated: (i) low-frequency subtraction (LFS); (ii) spatial propagation of information between probing windows, that is, by exclusion of the dependencies of the effector variable on the terms 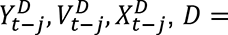 {*L*, *R*, *Up*, *Lo*}; in Eq. 2; (iii) Akaike-Information Criterion (AIC)-based model selection, that is, by pre-setting all the lag orders (*P*_0_, *P*_1_, *Q*_0_, *Q*_1_ and *M*_0_) to 1; and (iv) contemporaneous links. The four variants of the framework were applied to two imaging datasets in which we have strong expectations of GC outcomes based on existing literature: two-channel movies of (actin, Arp2/3) and (actin, cytoplasmic HaloTag).

Elimination of LFS had no effect on the prediction, in that the analyses for both data sets resulted in the same GC networks (Figure S8A). Thus, the LFS was not critical for the GC result, but it produced clearer cross-correlation curves and auto-correlation curves. In contrast, elimination of spatial propagation or AIC-based model selection, resulted in GC outcomes that were neither consistent with the outcomes generated by the complete modeling strategy nor were they consistent for common links between the two imaging datasets (actin, Arp2/3) and (actin, cytoplasmic HaloTag) (Figures S8B–C). For example, when we pre-specified all the lag orders (*P*_0_, *P*_1_, *Q*_0_, *Q*_1_ and *M*_0_) to be 1 without utilizing the AIC-based model selection, this analysis identified the GC from F-actin to edge velocity only in layer 1 for the dataset of (actin, Arp2/3) movies, whereas it identified the GC from F-actin to edge velocity only in layer 2 for the dataset of (actin, cytoplasmic HaloTag) movies (Figure S8C). This is a self-contradiction, demonstrating that over-simplified model orders can lead to wrong GC relations, and the model selection step is essential for accurate causal inference. The analysis without using the instantaneous GC identified no causal interactions between any of the variables (Figure S8D), suggesting that a proper model for lamellipodia dynamics imaged every 3 seconds needs to include contemporaneous predictors so that the model can capture fast interactions between actin and its regulators.

## SUPPLEMENTAL VIDEO LEGENDS

**Video S1. Fluorescence intensity of mNeonGreen-tagged actin (left) and endogenous Halo-tagged Arp3 (right) in a U2OS cell, related to** Figure 1

Imaging time interval is 3 sec/frame. Replay at 20 frames/second. Scale bar, 5 µm.

**Video S2. Protrusion/retraction vectors obtained by cell edge tracking, related to** Figure 1

Protrusion/retraction vectors connecting two consecutive cell boundaries (red arrows) are magnified (20-fold) for visualization. Scale bar, 5 µm.

**Video S3. Probing windows to extract molecular recruitments near the cell edge, related to** Figure 1

The size of probing windows is set to be ∼0.72 × 0.72 µm^2^. “w31” indicates the location of the 31^st^ window in the outmost band. Scale bar, 5 µm.

**Video S4. Reconstructed live cell movies displaying low-frequency (LF) oscillations and low-frequency subtracted (LFS) oscillations of actin assembly, related to** Figure 3

A sub-region of Video S1 (the 1^st^). Sampled actin activity shown in Figure 1H mapped back to the original pixel locations (the 2^nd^). LF oscillations (the 3^rd^) and LFS oscillations (the 4^th^) of actin activity mapped back to the original pixel locations. Protruding (red) and retracting (cyan) edge segments are annotated (white indicates unclassified). Past cell boundary traces 9 and 18 sec before are overlaid to assist visual inspection. The LFS oscillations co-fluctuate with the protrusion/retraction cycles, whereas the LS oscillation is unrelated to edge movement. Scale bar, 5 µm.

**Video S5. Spatial propagation patterns of actin intensity fluctuations in a fluorescence movie, related to** Figure 4A

Arrows in the four different directions indicate spatial propagation of actin fluctuations in a probing window to adjacent windows. Scale bar, 5 µm.

**Video S6. Spatial propagation patterns of cytoplasmic HaloTag fluctuations in a fluorescence movie, related to** Figure 4B

Arrows in the four different directions indicate spatial propagation of intensity fluctuations of a cytoplasmic probe in a probing window to adjacent windows. Scale bar, 5 µm.

**Video S7. Subcellular regions where Arp2/3 is Granger-causal for F-actin after accounting for the effect of edge velocity, related to** Figure 4C-4D

Magenta indicates the probing windows with a GC P-value < 0.05 superimposed on the Arp2/3 fluorescence movie. Scale bar, 5 µm.

**Video S8. Subcellular regions where F-actin is Granger-causal for Arp2/3 after accounting for the effect of edge velocity, related to** Figure 4C-4D

Magenta indicates the probing windows with a GC P-value < 0.05 superimposed on the actin fluorescence movie. Scale bar, 5 µm.

**Video S9. Subcellular regions where F-actin is Granger-causal for edge velocity after accounting for the effect of Arp2/3, related to** Figure 4C-4D

Magenta indicates the probing windows with a GC P-value < 0.05 superimposed on the actin fluorescence movie. Scale bar, 5 µm.

**Video S10. Subcellular regions where edge velocity is Granger-causal for F-actin after accounting for the effect of Arp2/3, related to** Figure 4C-4D

Magenta indicates the probing windows with a GC P-value <0.05 superimposed on the actin fluorescence movie. Scale bar, 5 µm.

**Video S11. Subcellular regions where Arp2/3 is Granger-causal for edge velocity after accounting for the effect of F-actin, related to** Figure 4C-4D

Magenta indicates the probing windows with a GC P-value < 0.05 superimposed on the Arp2/3 fluorescence movie. Scale bar, 5 µm.

**Video S12. Subcellular regions where edge velocity is Granger-causal for Arp2/3 after accounting for the effect of F-actin, related to** Figure 4C-4D

Magenta indicates the probing windows with a GC P-value < 0.05 superimposed on the Arp2/3 fluorescence movie. Scale bar, 5 µm.

**Video S13. Fluorescence intensity of exogenous SNAP-tagged mDia1 in a U2OS cell, related to STAR Methods**

Imaging time interval is 3 sec/frame. Replay at 20 frames/second. Scale bar, 5 µm.

**Video S14. Fluorescence intensity of endogenously labeled mNeonGreen2-mDia1 and SNAP-actin in a U2OS cell, related to** Figure 5C

Endogenous mNeonGreen2-mDia1 (left) and exogenous SNAP-actin (right). Imaging time interval is 3 sec/frame. Replay at 20 frames/second. Scale bar, 5 µm.

**Video S15. Fluorescence intensity of actin, wild-type and mutant VASP in a U2OS cell, related to** Figure 6A

mNeonGreen-actin, SNAP-tagged wild-type VASP (VASP^WT^), Halo-tagged S239D/T278E VASP mutant (VASP^MT^) that is actin-polymerization deficient, and VASP^WT^/VASP^MT^ merged from the left to right panel. Imaging time interval is 3 sec/frame. Replay at 20 frames/second. Scale bar, 5 µm.

## SUPPLEMENTAL TABLE LEGENDS

**Table S2. Wilcoxon signed rank P-values that determine Granger-causal pathways, related to** Figures 4–7, **S3, S6 and S7**

The P-values are from independent imaging experiments with (actin, Arp2/3), (actin, mDia1), (actin, VASP^WT^, VASP^MT^), (Arp2/3, VASP), and (actin, cytoplasmic HaloTag). The signed rank tests were implemented at different probing window layers at increasing distances from the cell edge from 0–0.7 µm (layer 1) to 2.2–2.9 µm (layer 4).

## Notes

### Competing Interest Statement

The authors have declared no competing interest.

