## Supplemental Figures and Tables for "Granger-causal inference of the lamellipodial actin regulator hierarchy by live cell imaging without perturbation"

### Figure S1

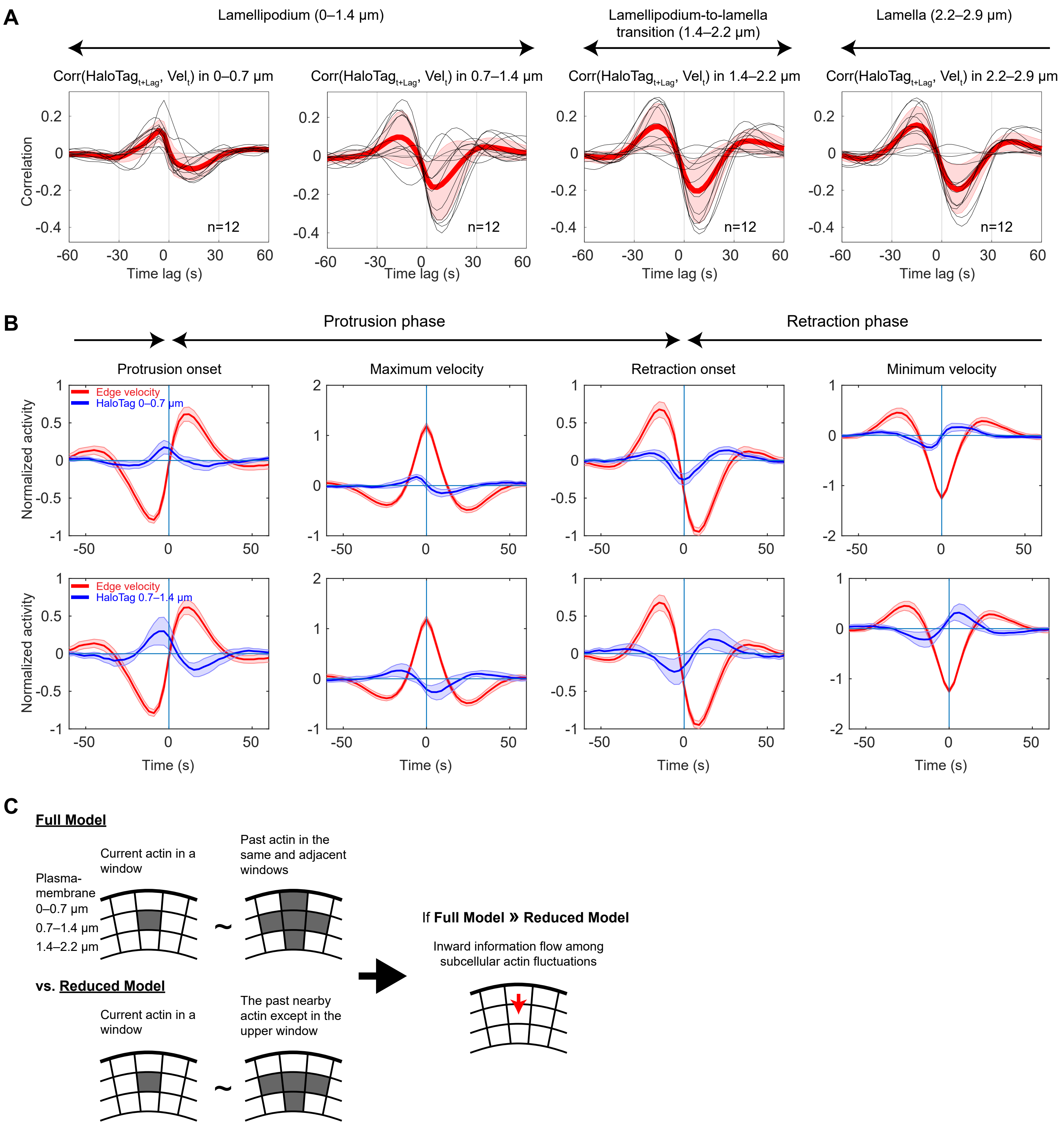


**Figure S1. Fluctuation patterns of cytoplasmic HaloTag intensities and illustration of the spatially propagating autoregressive (SPAR) model, related to Figures 3 and 4**
(A) Per-cell averaged cross-correlation curves (n = 12, black) of cytoplasmic HaloTag with the edge velocity. Red curves, averages over the cells. Cell-to-cell variability is shown by ±2 × SEM (shaded red bands). (B) Kinetic profiles of cytoplasmic HaloTag in the lamellipodia front (0–0.7 μm, top) and back (0.7–1.4 μm, bottom) during the cell edge motion events. Kinetic curves from different cells (n = 12) are averaged (solid lines), and shaded confidence bands indicate ±2 × SEM. (C) Illustration of the SPAR model to determine spatial propagation of molecular activities between adjacent probing windows.

### Figure S2

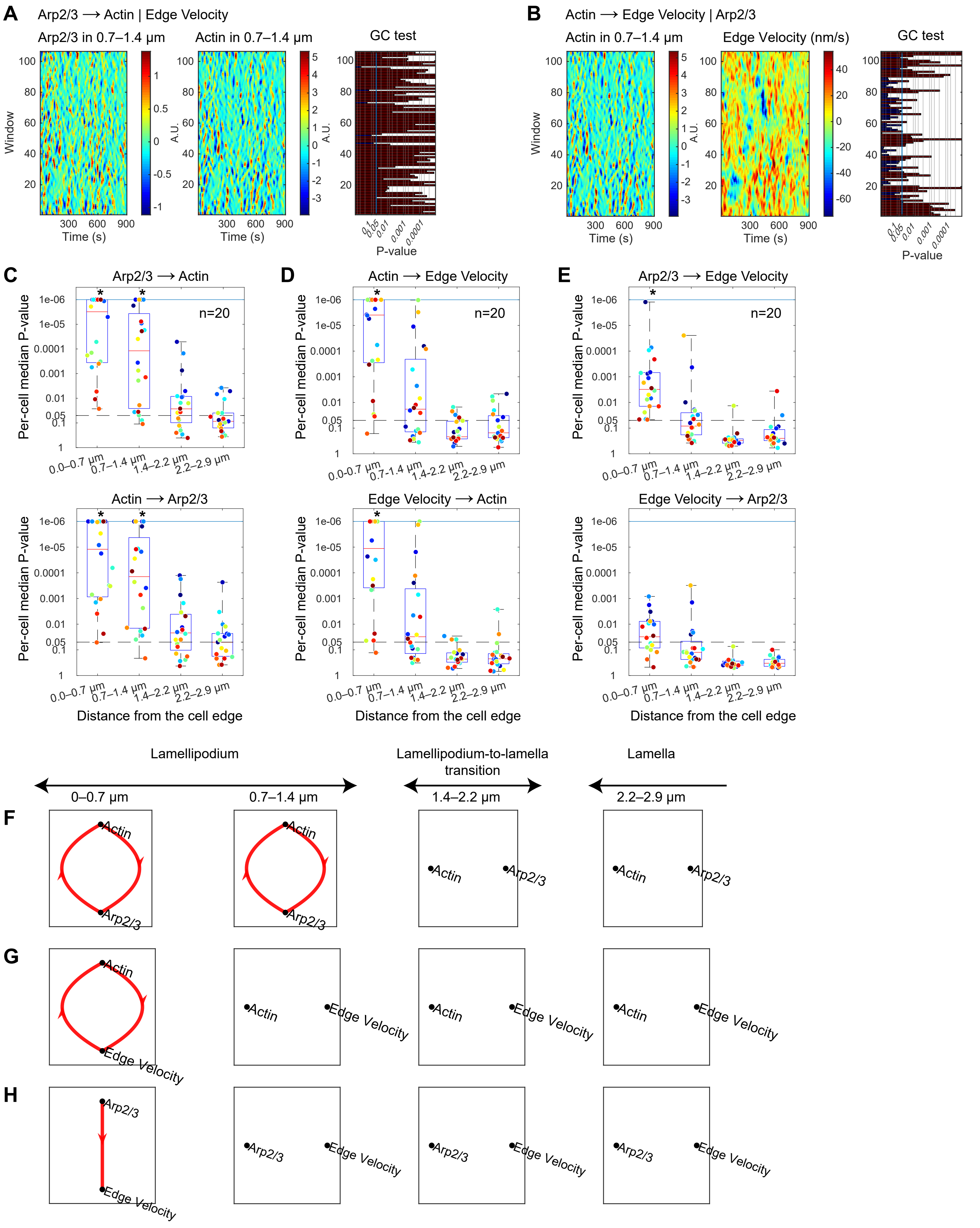
A
Arp2/3 ⟶ Actin | Edge Velocity
Arp2/3 in 0.7–1.4 μm
Actin in 0.7–1.4 μm
GC test
Window
Time (s)
A.U.
P-value
B
Actin ⟶ Edge Velocity | Arp2/3
Actin in 0.7–1.4 μm
Edge Velocity (nm/s)
GC test
C
Arp2/3 ⟶ Actin
Actin ⟶ Arp2/3
D
Actin ⟶ Edge Velocity
Edge Velocity ⟶ Actin
E
Arp2/3 ⟶ Edge Velocity
Edge Velocity ⟶ Arp2/3
n=20
Per-cell median P-value
0.0–0.7 μm
0.7–1.4 μm
1.4–2.2 μm
2.2–2.9 μm
Distance from the cell edge
Lamellipodium
Lamellipodium-to-lamella transition
Lamella
0–0.7 μm
0.7–1.4 μm
1.4–2.2 μm
2.2–2.9 μm
F
G
H
Actin
Arp2/3
Edge Velocity

**Figure S2. Granger-causality analyses of two-component systems embedded in a three-component observed system, related to Figure 4**

(A-B) Per-window P-values of the GC from Arp2/3 to actin (A) and from actin to the edge velocity (B) in the lamellipodia back (layer 0.7–1.4 μm) from the same cell as in Figure 4C-D. Red, significant P-values. (C-E) Boxplots of per-cell GC P-values (n = 20) when the GC is determined based on only two components among Arp2/3, actin and edge velocity, pretending that the other component is unobserved. The symbol (*) indicates that the per-cell P-values are significantly smaller than 0.05. (F-H) GC network diagrams of each of the embedded two-component systems consisting of Arp2/3, actin or edge velocity.

### Figure S3

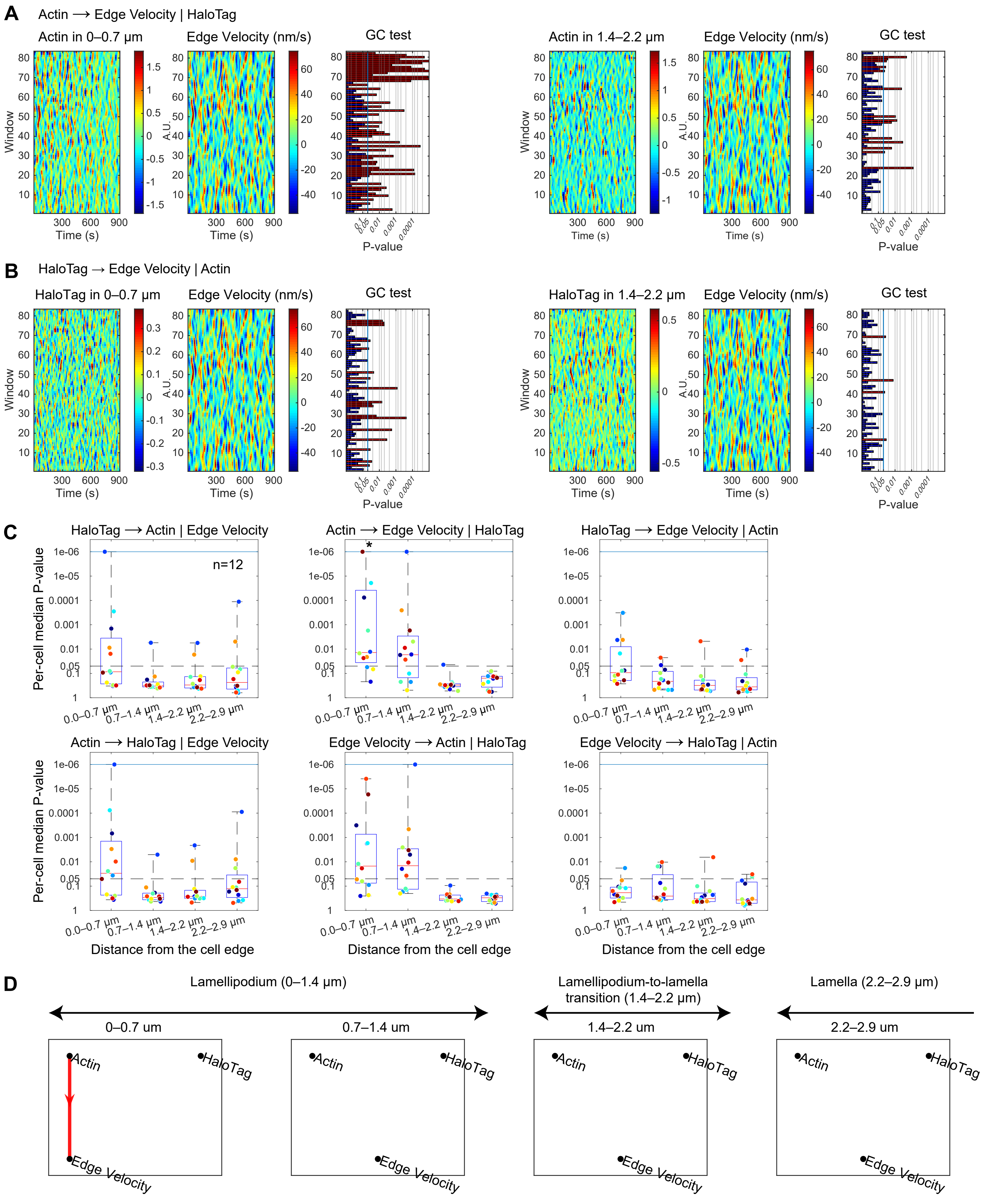

A
Actin ⟶ Edge Velocity | HaloTag
Actin in 0–0.7 μm
Edge Velocity (nm/s)
GC test
Actin in 1.4–2.2 μm
Edge Velocity (nm/s)
GC test
Window
Time (s)
A.U.
P-value
B
HaloTag → Edge Velocity | Actin
HaloTag in 0–0.7 μm
Edge Velocity (nm/s)
GC test
HaloTag in 1.4–2.2 μm
Edge Velocity (nm/s)
GC test
Window
Time (s)
A.U.
P-value
C
HaloTag ⟶ Actin | Edge Velocity
Actin ⟶ Edge Velocity | HaloTag
HaloTag ⟶ Edge Velocity | Actin
n=12
*
Per-cell median P-value
0.0–0.7 μm
0.7–1.4 μm
1.4–2.2 μm
2.2–2.9 μm
Actin ⟶ HaloTag | Edge Velocity
Edge Velocity ⟶ Actin | HaloTag
Edge Velocity ⟶ HaloTag | Actin
Distance from the cell edge
D
Lamellipodium (0–1.4 μm)
Lamellipodium-to-lamella transition (1.4–2.2 μm)
Lamella (2.2–2.9 μm)
0–0.7 um
0.7–1.4 um
1.4–2.2 um
2.2–2.9 um
Actin
HaloTag
Edge Velocity

**Figure S3. Cytoplasmic HaloTag does neither G-cause F-actin nor edge velocity, related to Figure 4**
(A-B) Activity maps of actin, cytoplasmic HaloTag and edge velocity, and their associated P-values of the GC from actin to edge velocity (A) and from HaloTag to edge velocity (B) in individual windows from the cell shown in Figure 4A-B. Red, significant P-values. (C) Boxplots of per-cell median P-values (n = 12) of Granger-causal relations between cytoplasmic HaloTag, actin and edge velocity. The symbol (*) indicates that the per-cell P-values are significantly smaller than 0.05. (D) GC pathway diagrams between cytoplasmic HaloTag, actin and edge velocity.

Figure S4

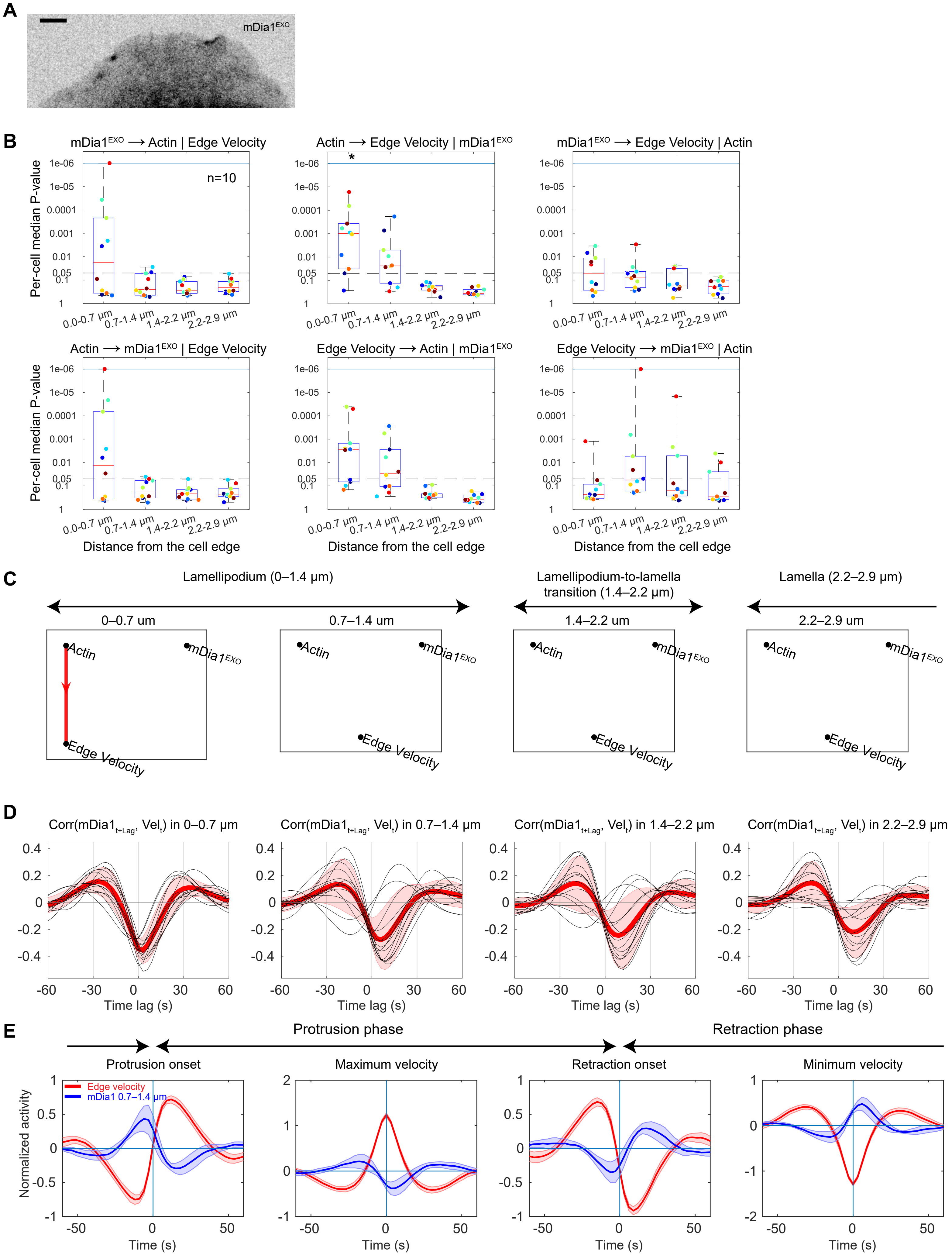
A
mDia1EXO
B
mDia1EXO ⟶ Actin | Edge Velocity
Actin ⟶ Edge Velocity | mDia1EXO
mDia1EXO ⟶ Edge Velocity | Actin
Actin ⟶ mDia1EXO | Edge Velocity
Edge Velocity ⟶ Actin | mDia1EXO
Edge Velocity ⟶ mDia1EXO | Actin
n=10
Per-cell median P-value
1e-06
1e-05
0.0001
0.001
0.01
0.05
0.1
1
0.0-0.7 μm
0.7-1.4 μm
1.4-2.2 μm
2.2-2.9 μm
Distance from the cell edge
C
Lamellipodium (0–1.4 μm)
Lamellipodium-to-lamella transition (1.4–2.2 μm)
Lamella (2.2–2.9 μm)
0–0.7 um
0.7–1.4 um
1.4–2.2 um
2.2–2.9 um
Actin
mDia1EXO
Edge Velocity
D
Corr(mDia1t+Lag, Velt) in 0–0.7 μm
Corr(mDia1t+Lag, Velt) in 0.7–1.4 μm
Corr(mDia1t+Lag, Velt) in 1.4–2.2 μm
Corr(mDia1t+Lag, Velt) in 2.2–2.9 μm
Time lag (s)
E
Protrusion phase
Retraction phase
Protrusion onset
Maximum velocity
Retraction onset
Minimum velocity
Edge velocity
mDia1 0.7–1.4 μm
Normalized activity
Time (s)

**Figure S4. Granger-causality analysis with exogenous mDia1 and fluctuation patterns of endogenous mDia1, related to Figure 5**

(A) Representative fluorescence image of exogenous SNAP-tagged mDia1 observed in a cell. (B) Boxplots of per-cell median P-values ($n = 10$) of Granger-causal relations between exogenous mDia1, actin and edge velocity. (C) GC pathway diagrams between exogenous mDia1, actin and edge velocity. (D) Per-cell averaged cross-correlation curves ($n = 14$, black) of endogenous mDia1 with the edge velocity. Red curves, averages over the cells. Cell-to-cell variability is shown by $\pm 2 \times$ SEM (shaded red bands). (E) Kinetic profiles of endogenous mDia1 during the cell edge motion events. Solid lines, averaged kinetic curves from different cells ($n = 14$); shaded bands, confidence bands ($\pm 2 \times$ SEM).

Figure S5

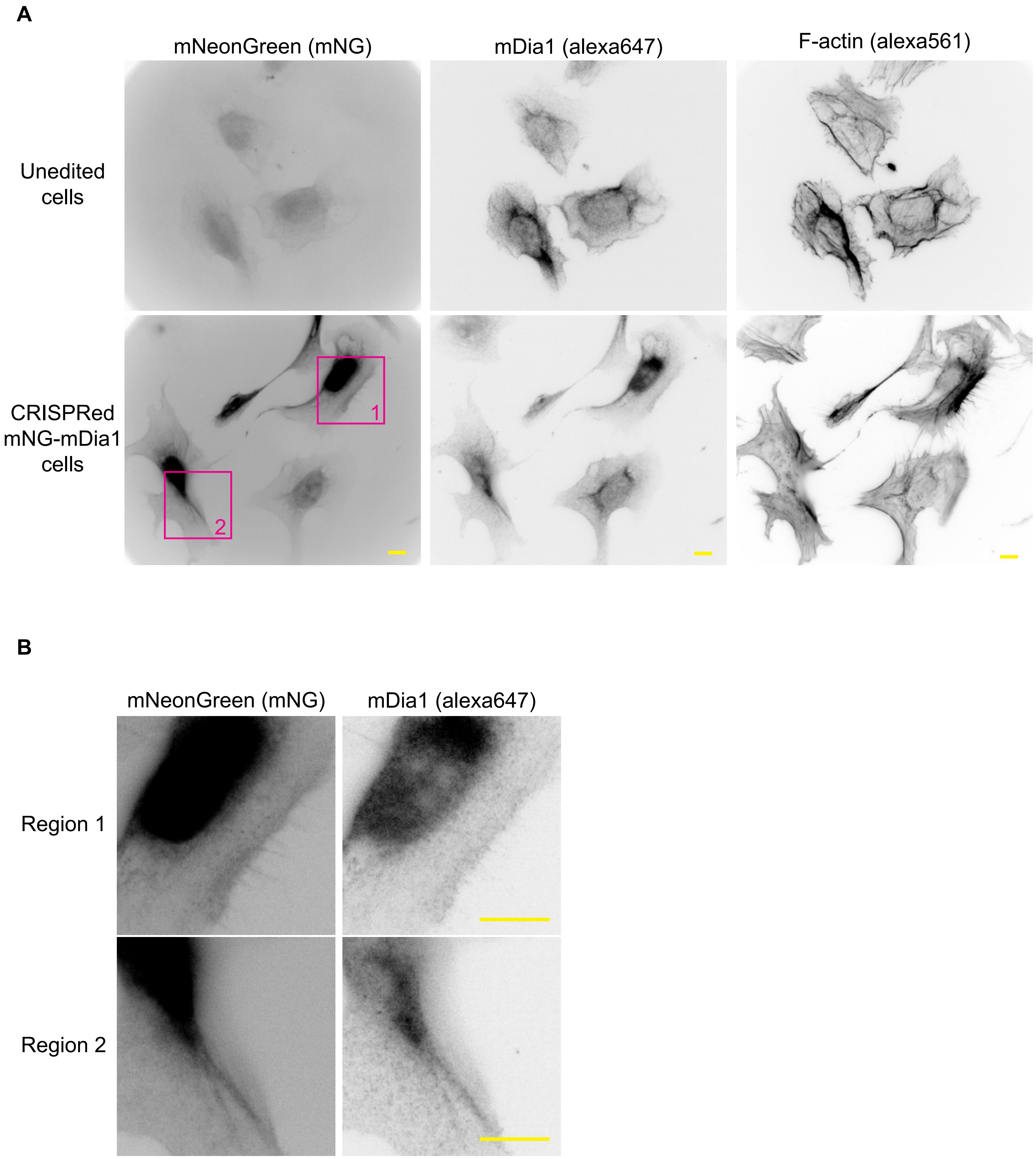


**Figure S5. Endogenously mNeonGreen211-tagged mDia1 does not alter mDia1 localization, related to Figure 5**
(A) Wild-type and endogenously mNeonGreen211-tagged mDia1 (mNG-mDia1) cells were seeded on fibronectin-coated glass coverslips and stained with anti-mDia1 antibodies and phalloidin. (B) Magnified images from the insets in (A) are shown. Scale bars are 10 μm.

**Figure S6**

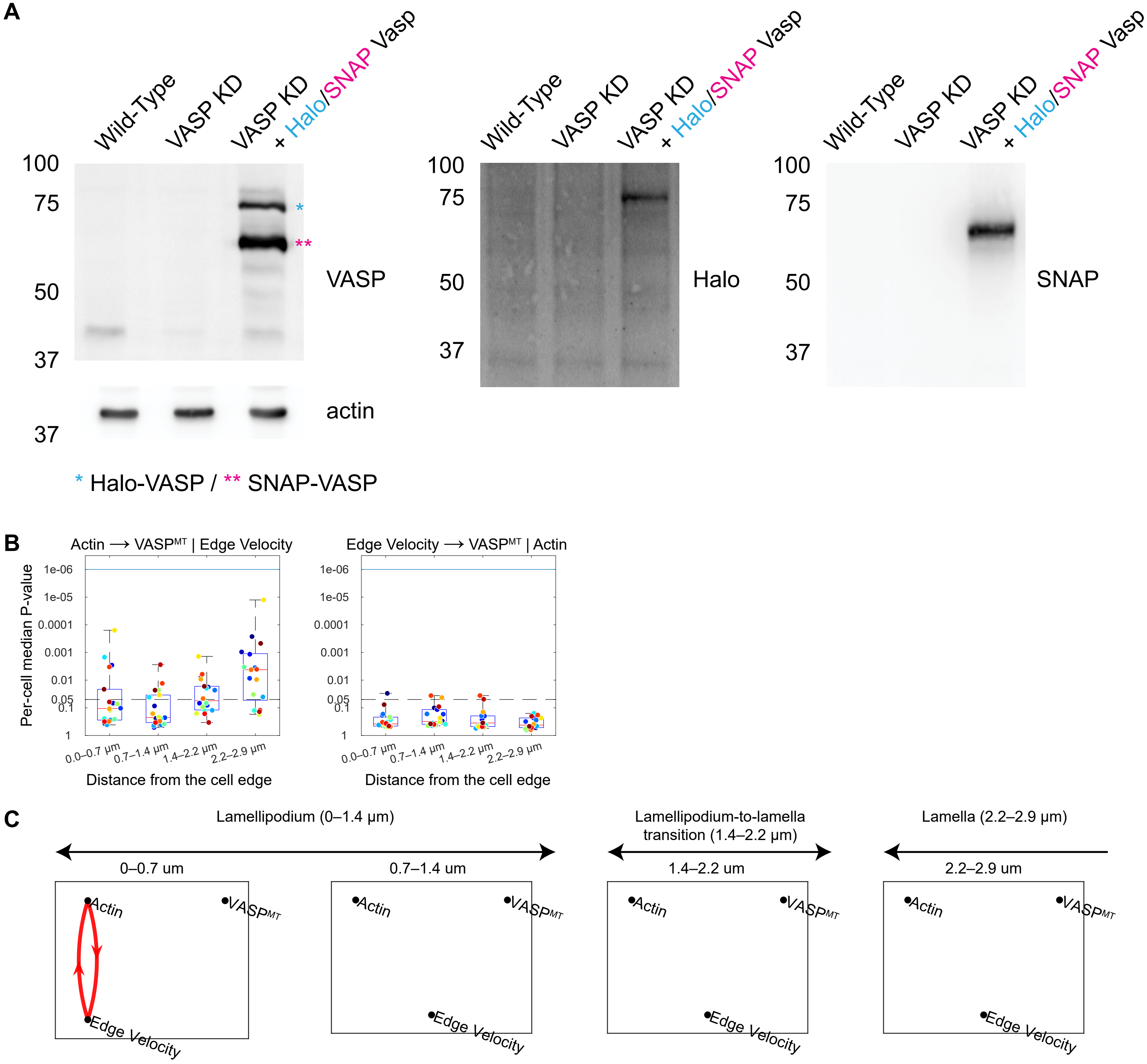


**Figure S6. Actin-polymerization deficient VASP does neither G-cause F-actin nor edge velocity, related to Figure 6**

(A) Western blot characterization of cells depleted for VASP (VASP KD) expressing exogeneous SNAP-tagged wild-type VASP (cyan asterisk) and Halo-tagged S239D/T278E mutant VASP (magenta asterisk). Wild-type cells served as a reference. SNAP-VASP and Halo-VASP were subsequently blotted with anti-SNAP and anti-Halo antibodies, respectively (right panels). (B) Boxplots of per-cell median P-values (n = 18) of Granger-causal relations between VASP mutant, actin and edge velocity. The remainder of boxplots is shown in Figure 6D-E. (C) GC pathway diagrams between VASP mutant, actin and edge velocity.

### Figure S7

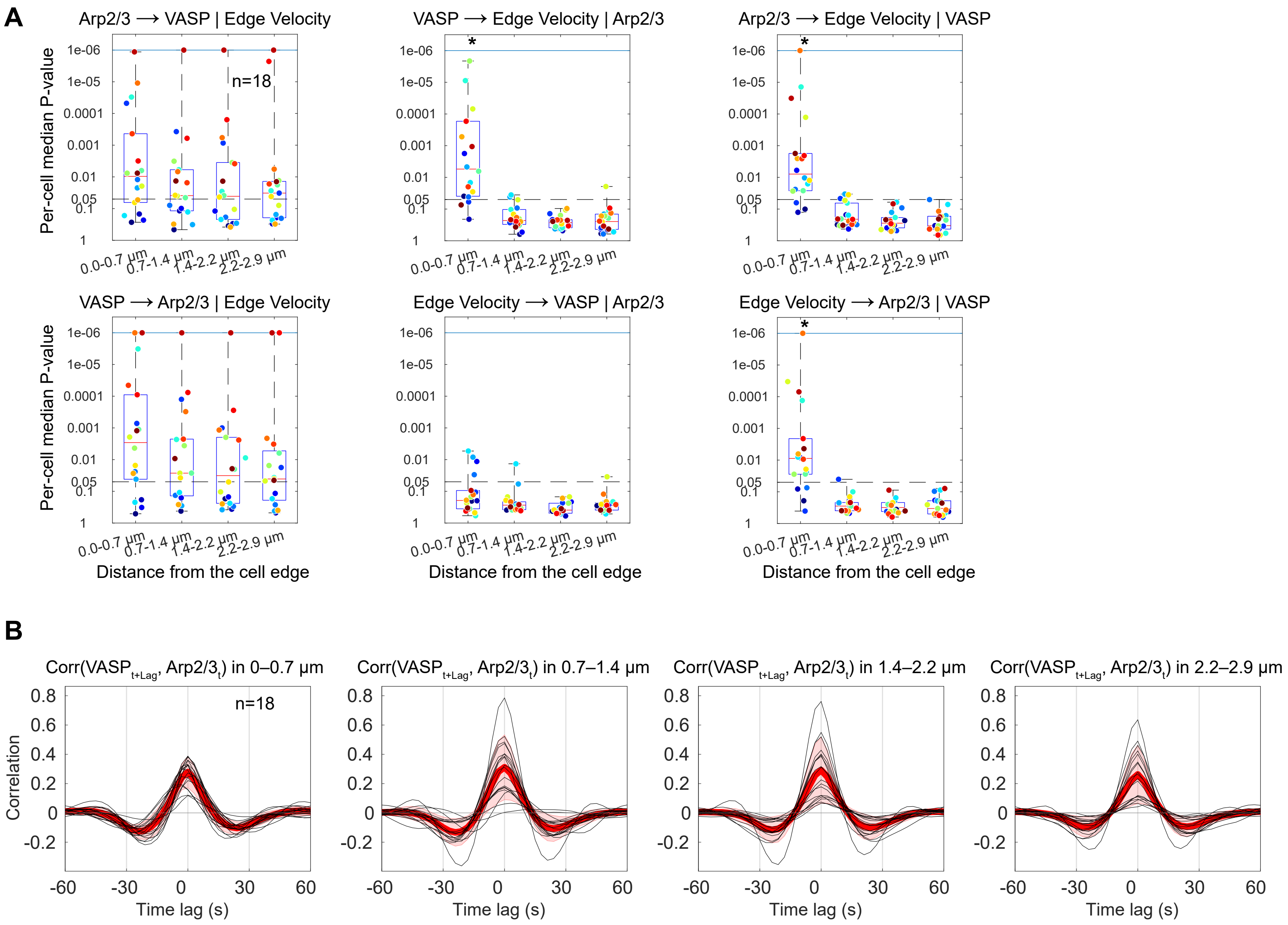


**Figure S7. Granger-causal hierarchy of actin regulators in the lamellipodia, related to Figures 4–7**
(A) Boxplots of per-cell median P-values (n = 18) of Granger-causal relations between Arp2/3, VASP and edge velocity at different distances from the cell edge. The symbol (*) indicates that the per-cell P-values are significantly smaller than 0.05 (Wilcoxon signed rank test). (B) Per-cell averaged cross-correlation curves (n = 18, black) of VASP with Arp2/3. Red curves, averages over the cells. Cell-to-cell variability is shown by ±2 × SEM (shaded red bands).

**Figure S8**

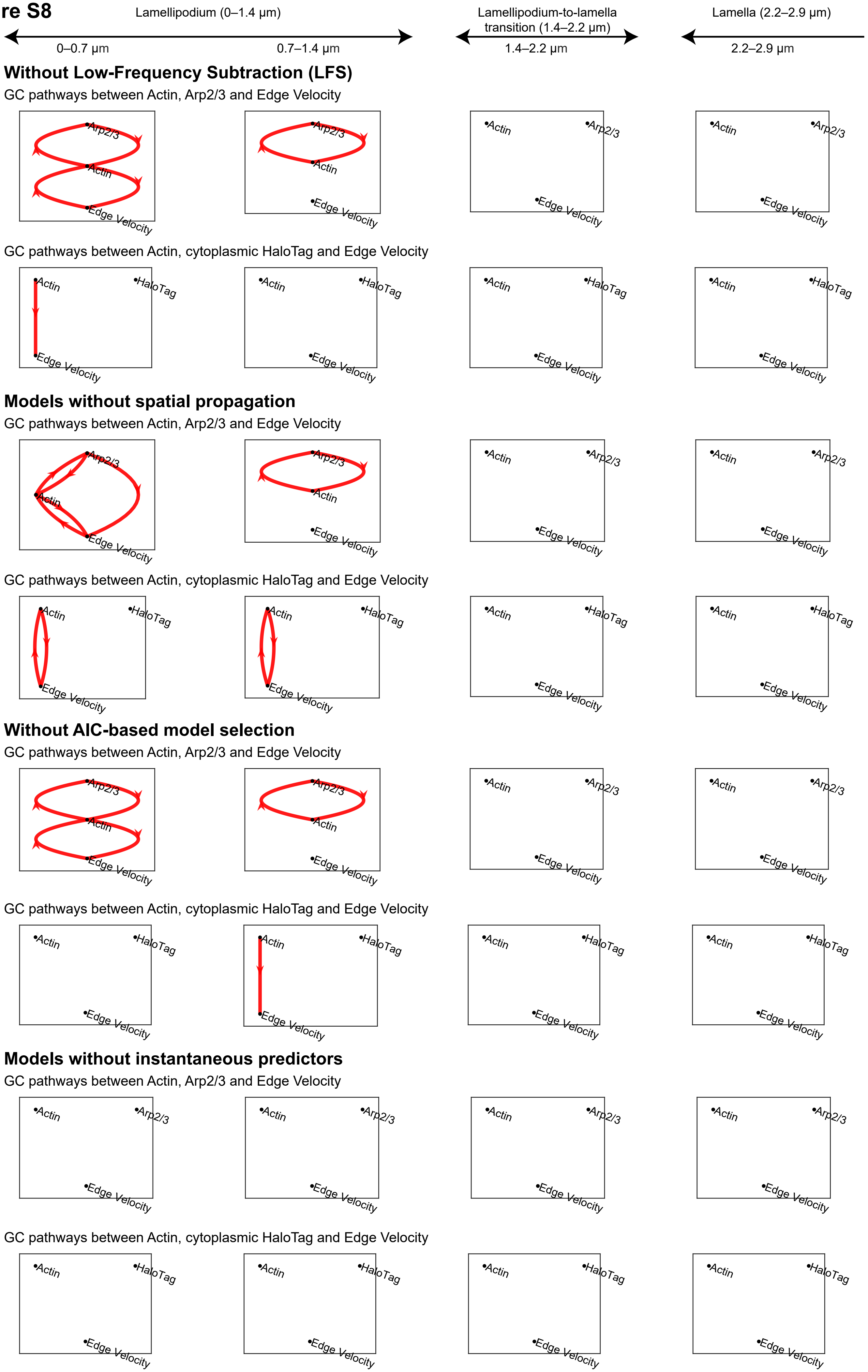

Lamellipodium (0–1.4 µm)
0–0.7 µm
0.7–1.4 µm
Lamellipodium-to-lamella transition (1.4–2.2 µm)
1.4–2.2 µm
Lamella (2.2–2.9 µm)
2.2–2.9 µm
A
Without Low-Frequency Subtraction (LFS)
GC pathways between Actin, Arp2/3 and Edge Velocity
GC pathways between Actin, cytoplasmic HaloTag and Edge Velocity
B
Models without spatial propagation
GC pathways between Actin, Arp2/3 and Edge Velocity
GC pathways between Actin, cytoplasmic HaloTag and Edge Velocity
C
Without AIC-based model selection
GC pathways between Actin, Arp2/3 and Edge Velocity
GC pathways between Actin, cytoplasmic HaloTag and Edge Velocity
D
Models without instantaneous predictors
GC pathways between Actin, Arp2/3 and Edge Velocity
GC pathways between Actin, cytoplasmic HaloTag and Edge Velocity
Actin
Arp2/3
HaloTag
Edge Velocity

**Figure S8. Granger-causality pathway outcomes subject to eliminating key modeling strategies, related to Figure 2**

GC pathway outcomes resulting from simplifications of the modeling strategy: (A) elimination of low-frequency subtraction (LFS); (B) elimination of spatial propagation; (C) elimination of Akaike-Information Criterion (AIC) based model selection; and (D) elimination of influences by contemporaneous variables. Shown are the GC pathway diagrams for the image data set (actin, Arp2/3) and (actin, cytoplasmic HaloTag).

### Supplemental Table

#### Table S1. List of oligonucleotides used in this study, related to STAR Methods

| Cloning | |
|---|---|
| Primer Name | Sequence 5' - 3' |
| mNeonGreen-18_fwd | atggcccgcctggctgaccgccgctagcgctaactagtGCCACCATGGTGAGCAAGGGCG |
| mNeonGreen-18_rev | gctctgaaagtacagatcctcagtggttggctcCTTGTACAGCTCGTCCATGCCCATCAC |
| Snaptag Actin Vector For | GCAAGCCTGGGCTGGGTTCCGGACTCAGATCTGGCAGCG |
| Snaptag Actin Vector Rev | CGCTTCATTTCGCAGTCTTTGTCCATGGTGGCACTAGTTAGCGCTAGCG |
| Snaptag Actin Insert For | CGCTAGCGCTAACTAGTGCCACCATGGACAAAGACTGCGAAATGAAGCG |
| Snaptag Actin Insert Rev | CGCTGCCAGATCTGAGTCCGGAACCCAGCCCAGGCTTGC |
| Halotag Actin Vector For | CGACGCTCGAGATTTCCGGCTCCGGACTCAGATCTGGCAGCG |
| Halotag Actin Vector Rev | GGAAAGCCAGTACCGATTTCTGCCATGGTGGCACTAGTTAGCGCTAGCG |
| Halotag Actin Insert For | CGCTAGCGCTAACTAGTGCCACCATGGCAGAAATCGGTACTGGCTTTCC |
| Halotag Actin Insert Rev | CGCTGCCAGATCTGAGTCCGGAGCCGGAAATCTCGAGCGTCG |
| SNAP-18_fwd | ggatgcgctgcctccaccgctgccagatctgagtccggaACCCAGCCCAGGCTTGCCCAG |
| SNAP-18_rev | gctctgaaagtacagatcctcagtggttggctcACCCAGCCCAGGCTTGCCCAGT |
| SpeI_Halo-18_fwd | attaactagtgccaccatggcagaaatcggtactggctttcc |
| XhoI_Halo-stop_rev | attctcgagTTACATAATTACACACTTTGTCTTTGACTTCTTTTTCTTC |
| Halo-18_fwd | gctgaccgccgctagcgctaactagtgccaccATGGCCACCATGGCAGAAATCGGTACTG |
| Halo-18_rev | tgatgcagagcccccaccgctgccgctccgaagaccggaCGACAGCCAGCGCGCGATCTC |
| linker_extender_rev | agatcttggccggccggcgatcgcgttatcGCTCTGAAAGTACAGATCCTCAGTGGTTGG |
| linker_extender_fwd | aaccactgaggatctgtactttcagagcgataacgcgaTCGCCGGCCGGCCAAGATCTTC |
| 18-actin_fwd | TGGGCTGGGTTCCGGACTCAGATCTGGCAGCG |
| 18-actin_rev | CAGTCTTTGTCCATGGTGGCACTAGTTAGCGCTAG |
| 18-mDia1_fwd | taacgcgatcgccggccggccaagatcttcGAGCCGTCCGGCGGGGGCCT |
| 18-mDia1_rev | ttttgtaatccagaggttgattgttccagacgcgtTTAGCTTGCACGGCCAACCAGCTCC |
| 18-VASP_fwd | aaccactgaggatctgtactttcagagcgataacgcgaTCGCCGGCCGGCCAAGATCTTC |
| 18-VASP_rev | ttttgtaatccagaggttgattgttccagacgcgtTCAAGGAGAACCCCGCTTCCTCAGC |
| ARP3_F1 | caccgttggagtcatgtcgtaaaat |
| ARP3_R1 | aaacattttacgacatgactccaac |
| ARP3_F2 | caccgggattgtgacgacaaatgct |
| ARP3_R2 | aaacagcatttgtcgtcacaatccc |
| shVASP_TRCN0000369620_fwd | CCGGAGGAATTGCAGAAAGTGAAAGCTCGAGCTTTCACTTTCTGCAATTCCTTTTTG |
| shVASP_TRCN0000369620_rev | AATTCAAAAAGGAATTGCAGAAAGTGAAAGCTCGAGCTTTCACTTTCTGCAATTCCT |
| 21-linker | GAGAACCACTGAGGATCTGTACTTTCAGAGCGATAACGCGATCGCCGGCCGGCCAAGATCTTC |

| Mutagenesis | |
|---|---|
| Primer Name | Sequence 5' - 3' |
| VASP_S235D_Fwd | GAcaaggaggaggcctctggggg |
| VASP_S235_Rev | cactttcctgagtttggctccagcaatgg |
| VASP_T273E_Fwd | GAacaggttggggagaagccccccaa |
| VASP_T273_Rev | ggcttttcttctccgggccagcatg |

| gBlock | |
|---|---|
| gBlocks | Sequence 5' - 3' |

| | |
|---|---|
| Kozak-ATG-mNG2-1.10 | actagtacttaaccatggtcgactggatccggtaccgaattcggtaccaccggtgcgccaccATGGTGTCAAAAGG<br>TGAGGAAGACAATATGGCGAGTCTTCCCGCGACGCATGAGTTGCATATATTTGGTTC<br>CATCAACGGCGTTGACTTCGATATGGTAGGTCAAGGGACAGGGAACCCAAATGACG<br>GGTATGAGGAACTCAACCTCAAGTCAACGAAAGGTGACTTGCAGTTCAGTCCGTGGA<br>TCTTGGTTCCTCACATCGGTTACGGCTTCCACCAATATCTTCCGTATCCCGACGGCA<br>TGTCACCCTTCCAAGCGGCGATGGTAGACGGAAGTGGTTACCAAGTCCACAGAACAA<br>TGCAATTCGAGGACGGTGCTAGCTTGACGGTCAATTATAGGTATACCTACGAGGGGA<br>GTCATATAAAGGGCGAGGCGCAAGTCATGGGAACTGGATTTCCGGCAGACGGACCA<br>GTAATGACCAATACCCTTACAGCTGCTGATTGGTGCATGAGCAAAAAGACTTATCCAA<br>ACGACAAGACAATCATTTCCACATTTAAGTGGTCATACACTACTGTCAACGGGAAACG<br>GTACCGATCAACAGCGAGGACCACCTACACTTTTGCTAAACCGATGGCAGCTAATTA<br>CCTCAAAAATCAGCCTATGTATGTATTTCGCAAAACCGAACTTAAGCATTCCATGgaatt<br>cgcggccgcactcgagatatctaacgcgttaatactagtatat |
| 4tandem_mNG2 with Linkers (mNG2_11x4) | ggtctcATGggatccACAGAGCTGAACTTCAAGGAGTGGCAAAAAGCATTCACAGATATGA<br>TGGGCAGCGGGTCCGGAACGGAACTTAACTTCAAAGAATGGCAAAAGGCATTTACG<br>GACATGATGGGGAGCGGATCAGGCACCGAGCTTAATTTTAAGGAATGGCAAAAAGCT<br>TTCACGGATATGATGGGATCAGGCAGCGGCACTGAATTGAATTTCAAGGAGTGGCAG<br>AAAGCTTTTACTGATATGATGTCAGGACTTCGGAGCGGTAGTGGAGGTGGGTCAGCT<br>TCCGGGGGTAGTGGGAGTagatctgagacc |
